# From CSP7 to CSP5: Development of a soluble, truncated congener for the treatment of obstructive and restrictive lung diseases

**DOI:** 10.64898/2026.11.13.766540

**Authors:** Akarsha Balnadupete, Kirana Mugaranja, Rashmi Shetty, Shalini Venkatesh, Ashoka Kumar, Joshua K Kleam, Hua Tang, Jay Peters, Nagarjun Venkata Konduru, Sreerama Shetty

## Abstract

Chronic obstructive and restrictive lung diseases (CLDs), represent a substantial worldwide health burden, with curative therapies remaining largely unavailable. Studies from our group and others have identified caveolin-1 (Cav1)-mediated signaling as a critical regulator of cellular processes implicated in the pathogenesis and progression of CLDs, including chronic obstructive pulmonary disease (COPD), idiopathic pulmonary fibrosis (IPF), silicosis and other progressive pulmonary fibrosis (PF) conditions. We have previously demonstrated that Cav1 scaffolding domain peptides (CSP and CSP7), attenuate lung injury and PF across multiple preclinical mouse models and exhibit protective and anti-fibrotic effects in COPD and IPF tissues ex vivo. The mechanism involves concurrent suppression of p53 and its downstream signaling in lung (airway and alveolar) epithelial cells by competitively counteracting excess Cav1, while restoring baseline p53 levels in (myo)fibroblasts persisting in fibrotic lungs, and alternatively polarized alveolar macrophages (AMs) by complementing Cav1 function. The safety and tolerability of inhaled micronized CSP7 dry powder (aka LTI-03), have been evaluated in mice, rats, dogs, followed by a Phase 1a clinical trial assessing safety and pharmacokinetics in healthy volunteers (NCT04233814) and a Phase 1b trial in patients with IPF (NCT05954988). LTI-03 has subsequently advanced to a multicenter Phase 2 clinical trial in patients with IPF (NCT06968845). However, the FDA highlighted findings of minimal bronchiolar mucus cell metaplasia and hyperplasia in the 26-week rat inhalation study, raisings concerns regarding the suitability of poorly soluble CSP7 for long-term administration in patients with CLDs, particularly for COPD, a disease already characterized by goblet cell metaplasia and hyperplasia, mucus hypersecretion, impaired mucociliary clearance, and small airway obstruction. To address these limitations, we identified CSP5, a five amino acid truncated derivative of CSP7. Notably, the amorphous CSP5 attenuated p53 activation and apoptosis in preclinical models of tobacco smoke (TS)-, bleomycin, and silica-induced lung injury, demonstrating its potential to protect against epithelial injury across diverse lung disease models. Additionally, CSP5 enhanced surfactant protein C expression, consistent with alveolar epithelial cell homeostasis and regeneration, while suppressing Mucin 5Ac expression and attenuating distal lung injury in mice chronically exposed to TS. Furthermore, CSP5 promoted resolution of established PF in mouse models of bleomycin- or silica-induced fibrosis. CSP7, encompassing the CSP5 sequence, attenuates autophagic dysregulation, endoplasmic reticulum stress, reactive oxygen species, mitochondrial membrane potential abnormalities, and mitochondrial superoxide production, and profibrotic polarization of AMs in mice with PF. Collectively, these findings position CSP5 as a formidable next-generation peptide with preserved bioactive motif of CSP7 while offering improved physicochemical and potential advantages in formulation flexibility. The dominant features of CSP5 support its further development for chronic inhaled or systemic administration in the treatment of obstructive or fibrotic lung diseases.

## Introduction

Chronic lung diseases are major causes of morbidity and mortality characterized by persistent epithelial injury, defective tissue repair, cellular senescence, sustained inflammation, and progressive remodeling of the airways and lung parenchyma. Long-term exposure to toxicants including tobacco smoke, particulates, radiation, and certain biological agents, damages airway and alveolar epithelial cells and initiates aberrant compensative repair responses that ultimately compromise lung architecture and function. A key feature of this pathologic process is loss of progenitor type II alveolar epithelial (AT2) cells due to increased apoptosis and senescence with reduced proliferative capacity^1–3^. Chronic epithelial injury is accompanied by recruitment and pro-fibrotic polarization of monocyte-derived alveolar macrophages (Mo-AMs), release of inflammatory cytokines and chemokines, oxidative stress, dysregulated protease–antiprotease activity, and accumulation of senescence-associated secretory phenotype (SASP) factors^4,5^. Furthermore, these responses establish a self-sustaining microenvironment that promotes epithelial dysfunction, fibroblast activation, and progressive structural remodeling of both the airway and alveolar compartments.

The predominant anatomical compartment affected, and the nature of the subsequent remodeling response determines whether this shared injury-repair program manifests primarily as airway obstruction, alveolar destruction, pulmonary fibrosis (PF), or a combination of these outcomes. In obstructive airway diseases such as COPD, chronic epithelial injury and inflammation drive goblet-cell hyperplasia and hypertrophy, mucus hypersecretion (MH), epithelial thickening, impaired mucus clearance, and small-airway narrowing, resulting in progressive airflow limitation. In diseases characterized by progressive fibrotic remodeling of the distal lung, such as PF, loss and senescence of AT2 cells impair alveolar regeneration, while persistent inflammatory and profibrotic signaling promotes activation and accumulation of fibrotic lung (myo)fibroblasts (fLfs). The activated fLfs demonstrate increased proliferation, migration and invasiveness, resistance to apoptosis, and excessive deposition of extracellular matrix (ECM). This leads to significant destruction of alveolar and interstitial architecture. Consequently, alveolar wall destruction and loss of elastic recoil can produce emphysema. Independent of the predominant obstructive or fibrotic manifestation, these chronic lung diseases (CLDs) share underlying core mechanisms of epithelial injury, AT2-cell senescence and apoptosis, inflammation and macrophage dysregulation, fibroblast differentiation and activation, oxidative stress, and impaired epithelial regeneration that collectively drive progressive loss of lung function^6–8^.

The considerable clinical burden and lack of curative therapies underscore the need to target the shared disease-driving mechanisms. Current therapies largely slow progression or control symptoms rather than restoring damaged lung architecture. In progressive PF, including idiopathic PF (IPF), approved antifibrotic therapies such as nintedanib, pirfenidone, and nerandomilast slow lung function decline but do not markedly prevent or reverse PF^9–12^. Likewise, current treatments for COPD predominantly improve symptoms, reduce exacerbations, and control inflammation without eliminating the underlying maladaptive injury-repair or reversing aberrant airway remodeling. These limitations support the identification and therapeutic targeting of common molecular pathways connecting persistent epithelial injury, cellular senescence, inflammatory signaling, fibroblast activation, and defective alveolar regeneration as a therapeutic strategy to potentially reverse the progressive remodeling that underlies CLD^4,6,7^.

Caveolae are cholesterol and sphingolipids rich lipid rafts that form small omega-shaped plasma membrane invaginations. Caveolae structure and function are maintained by coat proteins (caveolin1-3) and adaptor proteins (cavin1-4) and are involved in cell signaling, membrane transport, mechanosensing and transduction. Caveolin-1 (Cav1), a key structural and signaling protein of caveolar membranes, has long been recognized as a critical regulator of diverse protein- and lipid-mediated signaling pathways involved in numerous pathophysiological processes^13–18^. Caveolae and Cav1 are widely distributed in airway and alveolar epithelium, airway and pulmonary artery smooth muscle, AMs and fibroblasts, indicating their pathophysiological alterations associated with diverse types of lung diseases. Cav1 interacts with a variety of signaling molecules and particularly, regulates MAPK, NF-κB, TGFβ/Smad, and eNOS/NO signaling pathways in lung diseases^19^. In addition, Cav1 is involved in Ca^2+^ induced signaling and the hyper-responsiveness of airway smooth muscle^20^. Collectively, these findings identify Cav1 as a potential therapeutic target across diverse CLDs. Our group developed CSP7, a seven-amino-acid peptide derived from the Cav1 scaffolding domain, as a therapeutic strategy targeting Cav1-mediated pathological signaling in CLDs. As demonstrated in our recent reports^4,6,21,22^ and corroborated by others^23–26^, CSP7 has shown therapeutic efficacy across multiple preclinical models of chronic lung injury and progressive PF, including chronic tobacco smoke-induced lung injury (TSLI), bleomycin-induced PF (BLM-PF), constitutively active TGF-β1 adenovirus-induced PF (Ad-TGF-β1-PF), and silica-induced PF (sPF). CSP7 has been administered systemically by intraperitoneal (*IP*) injection following solubilization in DMSO-containing buffer, or directly to the lungs by nebulization (*Neb*) following solubilization in alkaline buffer with lactose as a carrier ^4,21^. It has also been delivered as a dry powder inhalation (*DPI*) formulation following micronization by air-jet milling (AJM)^21^ or formulation by thin-film freezing (TFF) ^27^. Across these complementary models and delivery approaches, CSP7 mitigated TSLI, resolved established PF, restored lung architecture, and significantly improved lung volume and pulmonary function.

CSP7, delivered systemically by *IP* injection or directly to the lungs by *Neb* or *DPI*, improves AT2-cell viability and telomere function, inhibits airway epithelial mucous cell (MC) metaplasia and MH, and reverses key pathological molecular changes in fLfs and Mo-AMs^4,6,21,27^. CSP7 solubilized in DMSO also demonstrates activity in human airway epithelial cells (AECs) and AT2 cells derived from COPD lungs, as well as in AT2 cells, fLfs, and Mo-AMs isolated from IPF lungs following *in vitro* treatment, with similar effects observed in *ex vivo*-treated human COPD and IPF lung tissues. Micronized CSP7, clinically developed as LTI-03, is currently being evaluated in a Phase 2 clinical trial in patients with IPF, following completion of a three-week Phase 1a study in healthy volunteers and a Phase 1b study in patients with IPF^28–33^.

However, poor solubility of CSP7 limits its suitability for systemic (intravenous or subcutaneous) and *Neb* delivery. Furthermore, long-term daily inhalation of poorly soluble CSP7 powder for the treatment of CLDs, such as COPD, may cause persistent airway irritation, raising potential concerns with prolonged exposure. Consistent with our thesis, the FDA noted minimal MC hyperplasia in the bronchioles following 24 weeks of CSP7 inhalation exposure in rats^34,35^. This treatment-related finding raises a particular concern for the long-term development of CSP7 for COPD, a disease already characterized by goblet-cell metaplasia/hyperplasia, MH, impaired mucociliary clearance (MCC), and small-airway obstruction. Although histologically minimal, chronic induction of this phenotype could potentially exacerbate an underlying pathologic feature of COPD and therefore represent an undesirable liability for a long-term inhaled therapy. This concern is further heightened by the absence of an established No-Observed-Adverse-Effect Level (NOAEL). Collectively, these limitations provide a strong rationale for developing an improved newer generation peptide that retains CSP7 therapeutic activity while overcoming its poor aqueous solubility and minimizing liabilities relevant to chronic treatment of COPD, particularly MC metaplasia/hyperplasia and MUC5AC (M5Ac)-driven MH. To address these limitations, we developed CSP5, a truncated five-amino-acid congener of CSP7 with markedly improved aqueous solubility. We directly compared aqueous-formulated CSP5 with the clinically advanced CSP7, which requires initial solubilization in DMSO followed by dilution in PBS, across multiple complementary *in vitro* and *in vivo* models of lung injury and existing PF. These studies were designed to determine whether amorphous CSP5 preserves the biological efficacy of CSP7 while providing physicochemical properties better suited for chronic pulmonary and systemic delivery.

## Materials and Methods

### Mouse models of Lung injury

Wild-type (WT) mice of C57BL/6J background were procured from Jaxon laboratory (Jax #000664). All *in vivo* experiments were conducted according to the protocol approved by the University of Texas Health Science Center at Tyler (UTHSCT) Animal Care and Use Committee (IACUC). Prior to the experiment, mice were acclimatized for a week in our vivarium facility located at UTHSCT campus.

### Synthesis and preparation of CSP5, CSP6 and CSP7 solutions for *in vitro* and *in vivo* interventions

The sequences of CSP7, CSP6, CSP5 and its deletion fragments were designed in our laboratory and commercially synthesized by GenScript. Following synthesis, the peptides were evaluated for their solubility in different solvents. Briefly, 20 mg of each peptide was dissolved separately in 500 µl of distilled H2O, Dulbecco’s phosphate-buffered saline (DPBS), regular saline (0.9% NaCl) or dimethyl sulfoxide (DMSO). The samples were mixed thoroughly and visually assessed for the formation of a clear solution to determine peptide solubility in each solvent.

### Treatment of BLM/Silica injured human AT2 cells with CSP5, CSP6 and CSP7 *in vitro*

Human ATII cells were procured from Cell Biologics (Chicago, IL) and cultured as mentioned in the earlier reports treated with silica (100 µg/ml) or BLM (40 µg/ml)^36,37^. Twenty-four hours later, injured cells treated with or without 20 µM of CSP5, CSP6, and CSP7 for additional 24h or 48h. The naïve cells, cells with silica or BLM injury left untreated or treated with above peptides were collected and used for further analysis.

### Preparation of TS extract and treatment of TS extract exposed ATII cells with CSP5, CSP6 and CSP7

TS extract was prepared using the research cigarettes 2R4F purchased from the Tobacco Health Research, University of Kentucky (Lexington, KY) as described in our earlier report^4^. Briefly, cigarettes were burned in a sidearm flask, and the generated smoke was bubbled into PBS at room temperature through an attached peristatic pump. Then the extract was filtered using a 0.2 um syringe filter and the absorbance of 1.0 measured at 230 nm was considered 100%. The ATII cells were cultured in 6-well plates and treated with TS extracts (0.4%) once the cells attain 60-70% confluency to induce the TS extract-based injury. After 24h, the cells were treated with or without CSP5, CSP6 and CSP7 (20 µM) in a fresh media and incubated for additional 48h before cells were harvested for protein and RNA extraction for further analysis.

### Development of mouse model of single dose (1X) BLM induced established PF (1X-BLM-PF)

WT mice were administered 1 U/kg BLM (Meitheal Pharmaceuticals, Inc) in 50 μl saline one time by intratracheal (*i.t.*) instillation to establish PF following BLM induced lung injury as we described elsewhere^21,27^. Mice exposed to same volume of vehicle (saline) were used as controls for comparison. Two weeks later when 1X-BLM-PF was florid and confirmed by pattern of post-BLM initial loss of body weight and subsequent stabilization *vs* saline controls, mice harboring BLM-PF were either left untreated (None) or subjected to daily *IP* injection of 1.5 mg/kg of CSP5, CSP6 and CSP7 for next 7 days from day 14 through day 21 as we described elsewhere^21,27^. The experiments were terminated by day 21 post-BLM and mice were subjected to micro-CT (μCT) scanning and pulmonary function testing (PFT). Later the mice were euthanized following the standard operating procedure set by IACUC, and lung tissues were harvested for further analyses.

### Development of TSLI induced PF *in vivo*

WT mice were exposed to daily 40 research grade cigarettes for 4h with 2h in between interval to have water and food using a mechanical smoking chamber (Teague Enterprises, Davis, CA) for 20 weeks to induce TSLI as we have described in our earlier publications^4^. Sixteen weeks after daily 4h exposure, mice with TSLI were treated with or without 1.5 mg/kg body weight CSP5 or CSP7 by *IP* injection. Naïve mice kept ambient air (AIR) were used as controls for comparison. At the end of the experiment *i.e.* 20 weeks after TSLI, naive mice and TSLI mice treated with or without CSP5 or CSP7 were subjected to μCT scanning and PFT as we reported earlier^4^. All mice were euthanized and lung tissues were harvested for further analysis.

### Development of Silica induced PF (sPF) *in vivo*

WT mice were exposed to 1.5 mg Silica in 100 μl saline by *i.t.* instillation as were reported earlier^36^. Fifty-four days later, mice with silica induced lung injury were left alone (None) or treated once daily with CSP5, CSP6 and CSP7 (1.5 mg/kg) by *IP* injection for 7 days from day 54 through day 60. Saline (vehicle) treated mice were used as controls. At the end of the experiments, all mice were euthanized, and lung tissues were harvested for further evaluation.

### Histology

Lung sections (5 mM) were deparaffinized and subjected to hematoxylin and eosin (H&E), Masson trichrome and picrosirius red staining as we described earlier to assess severity of lung injury and post-injury lung and airway remodeling as well as subsequent changes after treatment with CSP5 and CSP7. Images were captured simultaneously under identical circumstances in a blinded manner^27,36^.

### Periodic Acid Schiff’s (PAS) staining

PAS staining was performed according to the manufacture instructions (Abcam). Briefly, deparaffinized lung sections were treated with a 0.5% periodic acid solution for five minutes, washed in distilled water and later the sections were incubated with PAS stain for fifteen minutes with other reagents from the mucin staining kit^4^.

### μCT scanning, PFT and lung volume examination

At the end of the experiment, mice were subjected to μCT scanning; PFT and lung volume measurement were examined as described earlier^4,21,27^. Briefly, A 20-gauge intravenous (*IV*) canula was inserted through the vocal cords into the trachea after the mice were anesthetized. A flexiVent system (SCIREQ) using the “snapshot perturbation method” was utilized for PFT. This system assesses the elastance and compliance of the respiratory system, which shows the changes in the lung stiffness resulting from various treatments. As per the manufacture guidelines the tidal volume of 30 mL/kg at a breathing frequency of 150 breaths/min against 2-3 cm H2O positive end-expiratory pressure used in flexiVent, entire procedure was maintained under sedative using isoflurane throughout the PFT. Immediately after the PFT mice were subjected to CT imaging, using Locus μCT Scanner (General Electric, GE Healthcare). μCT scans with a resolution of 93 μm were carried out at full inspiration. Lung renderings obtained during full inspiration were used to determine lung volumes using Microview software.

### Immunoblot assay

AT2 cells and lung homogenate were sonicated with RIPA buffer (PMSF, protease inhibitor cocktail, and sodium orthovanadate) (Santa Cruz Biotech, #sc24948). Protein concentrations were quantified using Bicinchoninic acid (BCA) assay kit (Pierce, #23225). The equal amount of protein was loaded to the 4-20% gradient SDS-polyacrylamide gels (Genscript) and separated by electrophoresis (SDS-PAGE). Later, proteins were transferred to the nitrocellulose membrane (Bio-Rad, #1620112) and incubated with BSA (1%) at room temperature for 1h. Further, blots were incubated overnight with goat anti-collagen-1a1 (Col1) monoclonal antibody (1:1,000, Southern Biotech, #1440-01), rabbit anti-fibronectin1 (FN1) antibody (1:1,000, Molecular Innovations, #ASMFBN), and mouse anti-β-actin monoclonal antibody (1: 3,000, Cell signaling, #3700), anti-tenascin-C (TN-C) antibody (1:1,000, Cell signaling, #33352S), alpha-smooth muscle actin (a-SMA) antibody (1: 1,000, Cell signaling, #19245S), beclin-1 (1:500 santa cruz # sc-48381), ARG1(1:500 santa cruz # sc-271430), LC3B1(1:500 Cell signaling# 2775), SOCS3(1:500 santa cruz # sc-73045), P62(1:1000 Sigma# P0067) in Tris-Buffered Saline with Tween-20 (TBST) containing 1% BSA at 4° C. Later, membranes were washed using TBST for 1h, incubated with horseradish peroxidase (HRP)-conjugated anti-goat IgG (Jackson Lab, #705-035-003), and anti-rabbit IgG (Jackson Lab, #711-035-152) anti-mouse IgG (Jackson Lab, #715-035-150) secondary antibody diluted at 1:10,000 in TBST containing non-fat milk (5%) for 1h at room temperature. Later, washed membranes were later incubated with ECL Substrate (Pierce, #32106) and images were captured using a gel doc device (Syngene, Box-mini)^4,36,38^.

### Quantitative PCR (qPCR)

Total RNA was extracted from the lung homogenates and cell lysates using TRI reagent (MRC, #TR118). Equal amounts of isolated RNA were used for cDNA synthesis using iScript™ Reverse Transcription Supermix (Bio-Rad, #1708841). qRT-PCR was performed with iTaq™ Universal SYBR® Green Supermix (Bio-Rad, #1725121) using CFX96 Real-Time PCR System (Bio-Rad). The following primers Col1 (#qMmuCED0044222), FN1 (#qMmuCED0045687), α-SMA (qMmuCID0006375), TN-C (qMmuCED0047470) MUSC (#qMmuCED0039802), SP-C (qMmuCID0007880) and Gapdh (#qMmuCED0027497) were purchased from Bio-Rad^4,36,38^.

### Hydroxyproline assay

Lung homogenates were subjected to acid hydrolysis using 6N HCl for 24h at 120° C. Acid hydrolysate was evaporated and diluted with equal amount of Citrate/acetate buffer, transferred to a 96 well plate and added 100 μL of chloramine-T solution. Mixture was incubated in room temperature for 20 minutes followed by addition of 100 μl Ehrlich’s solutions and incubation of reaction mixture for 1h at 45° C. Absorbance was measured at 550 nm using a synergy H1 Hybrid reader (BioTek Instruments) using standard curve with known amount and total hydroxyproline content was calculated^21,27,36^.

### Soluble collagen (Sircol) assay

Lung homogenates were treated with 0.2 mg/ml pepsin (Acros Organics, #41707) in 0.5 M acetic acid for overnight at 4° C. Pepsin digests were centrifuged and clear supernatants were incubated with Sirius red dye for 30 minutes on the shaker at room temperature. Reaction mixture was centrifuged; pellets were rinsed with acid salt wash buffer and dissolved in alkaline reagent. Absorbance was measured at 550nm^21,27,36^.

### ROS, mitoSOX and mitochondrial membrane potential (MMP) assay

Reactive oxygen species (ROS) levels in ATII cells using cell-permeant reagent 2’,7’-dichlorofluorescin diacetate (DCFDA) (#ab113851) and mitochondrial (mt.) superoxide indicator, mitoSOX (#ab219943). We also analyzed MMP (ΔΨm) using the cell-permeant fluorescent dye, Tetramethylrhodamine ethyl ester perchlorate (TMRE).

### Statistical analysis

The statistical data analysis was using GraphPad Prism (9.0.0). Data represents as mean ± SD. Two-tailed unpaired Student’s t-test was used to assess statistical significance between two groups, whereas Turkey’s multiple comparison test and one-way ANOVA were used for multiple groups. The log-rank (Mantel-Cox) test was used to compute survival analysis. P < 0.05 was regarded as significant^4,21,27,36^.

## Results

### Improved Solubility of CSP5

The aqueous and solvent solubility profiles of acetate-form CSP5, CSP6, CSP7 and CSP were quantitatively assessed in ultrapure water, 0.9% NaCl solution, and dimethyl sulfoxide DMSO using a standardized solubility assay. Acetate-form peptides were first dissolved in the minimum volume of water required for complete solubilization (e.g. 1 ml of water for 1 mg of peptide with solubility of mg/ml). The resulting solution was then adjusted to physiological saline (0.9% NaCl) by adding an appropriate volume of concentrated NaCl solution. A peptide was classified as “freely soluble” if it dissolved rapidly to form a clear solution upon adding the solvent and vortexing or sonication. A peptide was classified as “insoluble” if turbidity or suspended material remained following vortexing or sonication. For quantitative assessment, peptide samples were initially evaluated at a concentration of approximately 20 mg/ml. Samples that did not dissolve at 20 mg/ml were sequentially diluted and re-evaluated at lower concentrations. CSP5 (TFTVT) demonstrated substantially improved aqueous solubility relative to CSP7 (FTTFTVT) or CSP6 (TTFTVT) or CSP. As shown in **Table 1**, the acetate-form of CSP5 was soluble in ultrapure water and 0.9% NaCl solution, whereas CSP, CSP7 and CSP6 were insoluble under the test conditions. However, all peptides were soluble in DMSO. Improved aqueous solubility facilitates preparation of inhalation formulations, including *Neb* solutions and *DPI* formulations, amenable for systemic (*IV* or subcutaneous) delivery, and may streamline manufacturing process while enhancing formulation feasibility for long-term therapeutic administration. Improved aqueous solubility enhances the pharmaceutical developability of the shorter peptide, CSP5, by facilitating the preparation of inhaled formulations and support CSP5 as a promising lead therapeutic candidate for treatment of COPD and related CLDs.

**Table 1:**
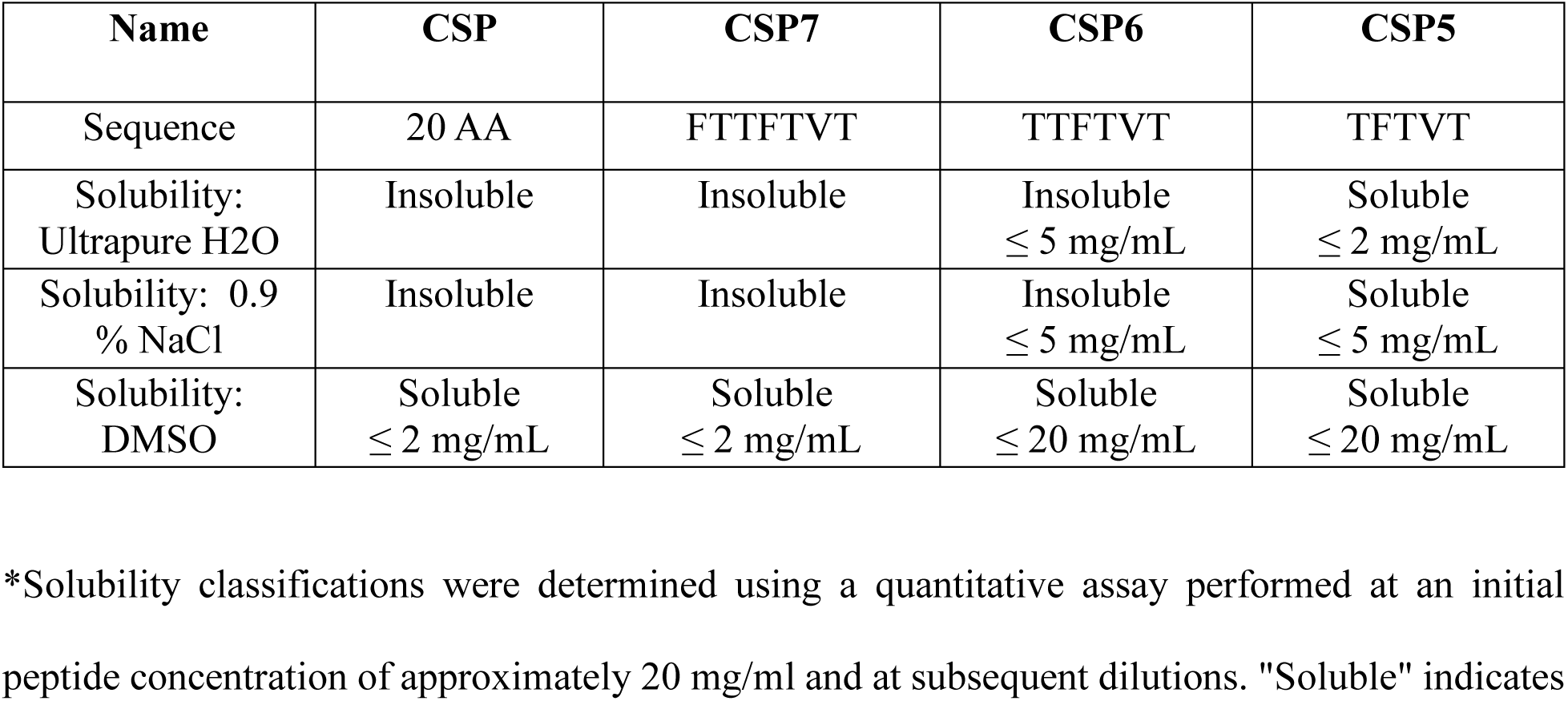
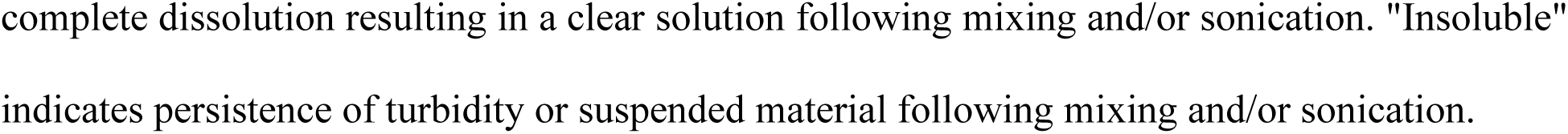
Solubility of CSP and its Derivatives.

### CSP7 deletion peptides protect human AT2 cells from BLM injury and resolve 1X-BLM-PF in WT mice

Human AT2 cells were exposed to BLM (40 µg/ml) for 24h, followed by treatment with CSP5, CSP6 and CSP7 for 48h. Total cellular protein was subsequently extracted and analyzed to assess the expression of p53, PAI-1, and surfactant protein-C (SP-C) expression as well as activated caspase-3 (Cl. Cas-3) levels as an indicator of apoptosis. As shown in **Figure 1A**, treatment of BLM-injured primary human AT2 cells with CSP5, CSP6 and CSP7 markedly reduced the expression of Cl. Cas-3, a key marker of apoptosis, compared to AT2 cells that received no subsequent treatment with peptides after BLM injury. Rescue of healthy AT2 cells after BLM injury by CSP7 and deletion peptides was further reflected by remarkable increase in SP-C protein and inhibition of p53 and PAI-1. These findings suggest that CSP5, CSP6 and CSP7 promote restoration of AT2 cell homeostasis by attenuating BLM-induced p53 and PAI-1 signaling associated with apoptosis.

**Figure 1.**
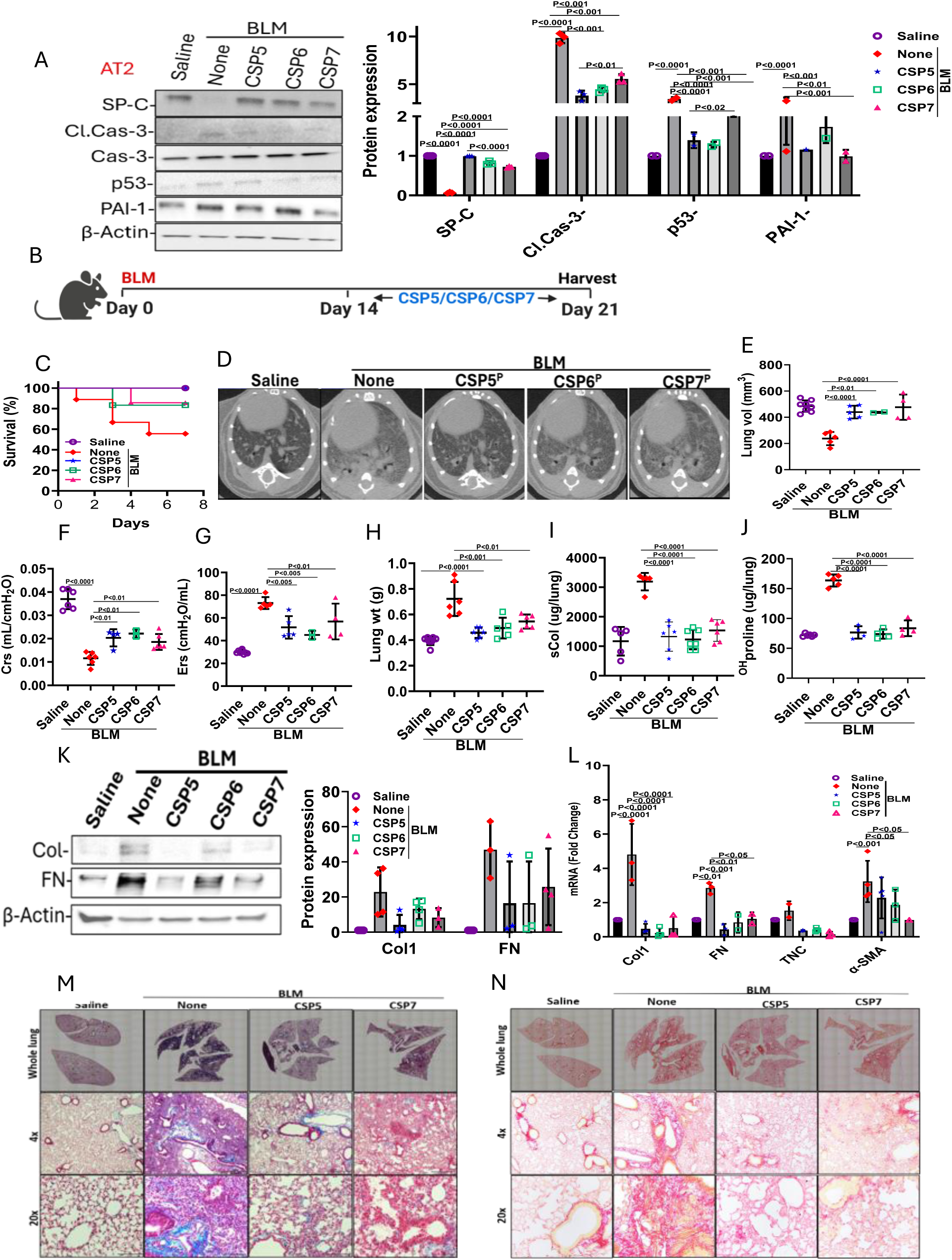
CSP7 deletion peptides inhibit BLM induced injury in human AT2 Cells and resolves BLM-PF in WT mice. (**A**) Human AT2 cells were exposed to BLM (40 mg/ml). After 24h, AT2 cells having BLM injury were treated with 10 mM CSP5, CSP6, and CSP7 for additional 48h. AT2 cell lysates were analyzed for SP-C, Cl. Cas-3, PAI-1 and b-actin by Western blotting. The bar graph represents densities of individual bands (n=3) after normalization with corresponding b-actin loading controls. (**B**) Schematic representation of experimental design. WT mice (n=5) were administered BLM by *i.t.* instillation to induce PF. Fourteen days later, mice with BLM-PF were treated with CSP5, CSP6, or CSP7 at approximately 1.5 mg/kg by daily IP injection for next seven days. (**C**) Survival (%), (**D**) μCT Images, (**E**) Lung volume after μCT rendition, (**F**) Lung compliance (Crs), (**G**) Lung elastance (Ers) and (**H**) Lung weights were measured. Whole lung homogenates were analyzed for (**I**) sCol, (**J**) ^OH^Proline, (**K**) Profibrotic marker proteins by Western blot. The bar graph represents densities of individual bands (n=3) after normalization with corresponding b-actin loading controls. (**L**) Total lung RNA was analyzed for profibrotic marker mRNA by qPCR. Lung sections were subjected to (**M**) Trichrome and (**N**) Picrosirius staining.

CSP5 and CSP6 recapitulate the anti-apoptotic effects of CSP7 in human AT2 cells. Multiple studies from our group have demonstrated that CSP7 attenuates established BLM-PF^21,22,27^. Therefore, to determine whether CSP7-derived deletion peptides, particularly CSP5, which exhibit greater water solubility, retain the anti-fibrotic activity of the parent peptides (CSP7 or CSP), we evaluated their therapeutic efficacy in WT mice with established BLM-PF. As shown in **Figure 1B**, mice were treated with CSP5, CSP6 and CSP7 at a dose of 1.5 mg/kg via *IP* injection daily for seven consecutive days from day 14-21. Experiments were terminated 24h after the last dosing of peptides. As shown in **Figure 1C**, CSP6- and CSP7-treated groups exhibited survival rates of approximately 80-90% compared to only 50% survival rate in untreated BLM-PF mice. In contrast, CSP5 treatment resulted in 100% survival in BLM-PF mice. The μCT images revealed minimal residual fibrotic scarring in BLM-PF mice treated with CSP5, CSP6 or CSP7 (**Figure 1D**). Further, μCT-derived lung volume measurements and SCIREQ-based lung function measurements demonstrated that of BLM-PF mice treated treatment with CSP7 or its deletion analogs significantly improved lung volume (**Figure 1E**) and compliance (**Figure 1F**), while reducing lung elastance (**Figure 1G**), thereby reversing the impairments observed in untreated BLM-PF mice.

Similarly, treatment of BLM-PF mice with CSP7 or its truncated derivatives significantly reduced gross lung weight (**Figure 1H**), soluble collagen (sCol) levels (**Figure 1I**), and total hydroxyproline (^OH^Proline) (**Figure 1J**) content in whole-lung homogenates, supporting their antifibrotic efficacy. Furthermore, immunoblot analysis of whole-lung homogenates revealed marked reduction in profibrogenic marker proteins such as Col1 or FN1 proteins following treatment of BLM-PF mice with CSP7 or its shorter derivatives (**Figure 1K**). These findings were further corroborated by significant reductions in the mRNA expression of key profibrotic markers, including Col1, FN1, TN-C, and a-SMA in 1X-BLM-PF mice treated with CSP5, CSP6, or CSP7 compared with untreated BLM-PF mice (**Figure 1L**). Consistent with the significant reductions in sCol, ^OH^Proline, and Col, FN1 and other ECM proteins and their mRNA levels in whole lung homogenates, histological evaluation of Masson’s trichrome-(**Figure 1M**) and picrosirius red-stained (**Figure 1N**) lung sections revealed remarkable reduction in interstitial deposition of Col and other ECM proteins in 1X-BLM-PF mice treated with CSP5, CSP6 and CSP7 peptides.

### CSP5 and CSP7 attenuates TS induced injury in human AT2 cells and mitigate TSLI in WT mice

We next evaluated the protective effects of CSP5 on TS-induced alveolar epithelial injury, wherein human AT2 cells were exposed to TS extract for 24h, followed by treatment with or without CSP5 or CSP7 (20 µM) for an additional 48h. Consistent with our observations following BLM-induced injury, exposure of AT2 cells to TS extract increased the activation of Cas-3 and PAI-1, suggesting enhanced apoptotic signaling and epithelial injury from TS exposure (**Figure 2A**). This was further evidenced by reduced expression of SP-C in AT2 cells exposed to TS extract. Conversely, treatment of TS extract-injured AT2 cells with CSP5 or CSP7 markedly attenuated AT2 cell injury reflected by restoration of SP-C expression and suppression of, an apoptotic marker, and PAI-1, a key mediator of p53-associated cellular senescence, compared with injured AT2 cells that were not treated with peptides. These findings suggest that CSP5 retains the cytoprotective effects of CSP7 and promotes restoration of AT2 cell homeostasis following TS-induced injury.

**Figure 2.**
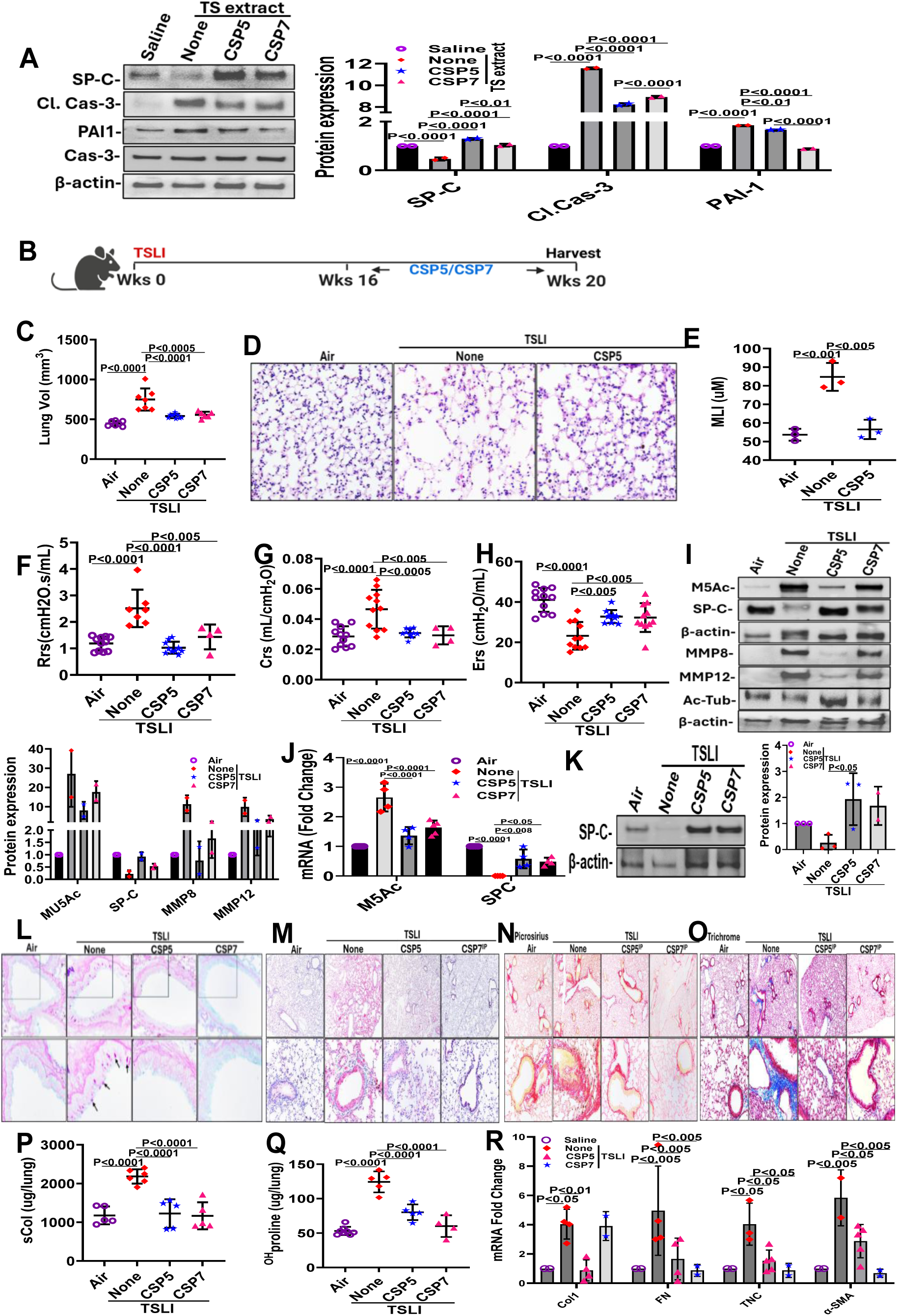
CSP5 and CSP7 ameliorate the TS induced injury in AT2 cells *in vitro* and mitigate airway and alveolar injury in WT TSLI mice. **(A)** AT2 cells were treated with TS extract (0.4%). One day later, AT2 cells exposed to TS extract were treated with or with 10 mM CSP5 or CSP7 for an additional 48h. AT2 cell lysates were analyzed for SP-C, Cl. Cas-3, PAI-1 and b-actin by Western blotting. **(B)** Schematic representation of experimental design. WT mice (n=5-10/gp) were kept in ambient Air or exposed to TS for 20 weeks to induce TSLI. After 16 weeks, mice having TSLI were left untreated (None) or treated with CSP5 or CSP7 (1.5 mg/kg/day) by IP injection for 5 days a week for the last 4 weeks. **(C)** Lung volume measured after μCT rendition and **(D)** H&E staining and **(E)** Mean linear intercept (MLI) of lung sections. (F) Lung resistance (Rrs) **(G)** lung compliance (Crs), and **(H)** Lung elastance (Ers) were measured. Whole lung homogenates were analyzed for **(I)** Expression M5Ac, SP-C, MMP-8, MMP-12 and acetylated tubulin (Ac-Tub) by Western blotting. The bar graph represents densities of individual bands (n=3) after normalization with corresponding b-actin level. **(J)** Total lung RNA was analyzed for M5Ac and SP-C mRNA by qPCR. **(K)** Lysates of AT2 cells isolated from TSLI mice treated as in Fig 2I immunoblotted for SP-C. The densities of individual bands (n=3) presented as bar graph after normalization with corresponding b-actin. Lung sections were subjected to (L) PAS staining to visualize mucin secretion, black Arrows indicate PAS-positive staining or (M) IHC for M5Ac or (**N**) Picrosirius and (**O**) Trichrome staining. Whole lung homogenates from of above treated mice were analyzed for (**P**) sCol and (**Q**) ^OH^Proline contents and (R) Profibrotic markers mRNA.

Our previous studies^4^ demonstrated that CSP7, administered via *IP* or delivered directly to the airways through *Neb* following solubilization in NaOH or as *DPI* after micronization prepared by AJM, improved lung function and attenuated airway and alveolar pathology in a mouse model of TSLI. In the present study, we compared the therapeutic efficacy of CSP5 and CSP7 in mitigating TSLI in WT mice. Exposure of WT mice to TS for 4h per day, 5 days per week, over 20 weeks (**Figure 2B**) resulted in a significant increase in lung volume compared with mice exposed to ambient-air, consistent with the development of COPD-like lung pathology. Importantly, treatment with either CSP5 or CSP7 during the final four weeks notably restored elevated lung volumes observed in TSLI toward those measured in ambient air-exposed controls (**Figure 2C**), consistent with reduced TS-induced alveolar injury. The attenuation of alveolar injury in CSP5-treated mice was further confirmed by histopathological evaluation of H&E-stained lung sections (**Figure 2D**) and quantitative assessment of the mean linear intercept (**Figure 2E**). These improvements in lung architecture were accompanied by improved lung function, as evidenced by reduced airway resistance (**Figure 2F**) and lung compliance (**Figure 2G**), along with increased lung elastance (**Figure 2H**) in CSP5- or CSP7-treated mice compared with untreated TSLI controls.

The protective effects of CSP5 against TS-induced alveolar and airway injury were further evaluated through molecular analyses of key markers associated with lung epithelial damage and remodeling. Consistent with the improvements in lung volume and PFT readouts, immunoblot analysis of whole-lung homogenates and subsequent densitometric quantification revealed that CSP5 treatment restored SP-C expression and acetylated tubulin (Ac-Tub), which was significantly reduced in untreated mice with TSLI, while concurrently suppressing the expression of MMP-8, MMP-12, and airway mucus-associated protein M5Ac. Notably, the effects of CSP5 on the expression of SP-C, M5Ac, Ac-Tub, MMP-8 and MMP-12 were more pronounced than those observed following CSP7 treatment in TSLI mice (**Figure 2I**), suggesting superior efficacy of CSP5 in mitigating TS-induced airway and alveolar injury. Consistent with the protein-level changes, qPCR analysis of lung homogenates confirmed that CSP5 treatment reduced M5Ac mRNA expression while restoring SP-C transcript levels (**Figure 2J**). Restoration of SP-C expression was further validated by immunoblotting of AT2 cells isolated from the TSLI mice treated with CSP5 or CSP7 and compared with TSLI controls (**Figure 2K**). To further investigate the effects of CSP5 and CSP7 on TS-induced airway MH and goblet-cell metaplasia, lung sections were subjected to periodic acid-Schiff (PAS) staining (**Figure 2L)** and IHC analysis using anti-M5Ac antibody (**Figure 2M**). Histological evaluation revealed markedly increased PAS-positive goblet cells and airway mucin accumulation in untreated TSLI mice, consistent with goblet-cell metaplasia and MH. Contrastingly, treatment with CSP5 and CSP7 substantially reduced goblet-cell accumulation and M5Ac-positive staining within the airway epithelium, corroborating the molecular findings and demonstrating attenuation of TS-induced airway mucus pathology.

Airway remodeling is a prominent pathological feature in patients with COPD, and emerging evidence suggest co-occurrence of PF and emphysema, collectively termed combined PF and emphysema (CPFE), representing a distinct clinical phenotype associated with substantial morbidity and mortality^39,40^. Given the potent anti-fibrotic effects of CSP5 and CSP7 demonstrated in 1X-BLM-PF mice, lung sections from TSLI-mice were subjected to picrosirius red staining to assess collagen deposition (**Figure 2N)** and Masson’s Trichrome (**Figure 2O)** staining to evaluate accumulation of collagen and other ECM proteins, indicative of fibrotic tissue remodeling. Histological analysis revealed increased peribronchial deposition of collagen and other ECM proteins in untreated TSLI mice. Notably, treatment with CSP5 and CSP7 peptides, markedly attenuated the peribronchial collagen deposition to near-baseline levels observed in ambient AIR-exposed controls. Furthermore, analysis of the whole-lung homogenates revealed significant increases in sCol (**Figure 2P**) and total ^OH^Proline (**Figure 2Q**) in untreated TSLI mice compared with ambient air-exposed controls. Consistent with the anti-fibrotic efficacy demonstrated in BLM-PF mice (**Figure 1**), treatment with CSP5 and CSP7 significantly reduced sCol and ^OH^Proline levels in the lungs of TSLI mice. These reductions were accompanied by significant downregulation of mRNA expression of key profibrotic markers, including Col1, FN1, TN-C, and a-SMA in CSP5-and CSP7-treated mice compared with untreated TSLI controls (**Figure 2R**), further supporting their potential to attenuate TSLI-driven fibrotic remodeling.

### CSP7 inhibits the silica induced ROS and restores MMP

Inhalation exposure to particulates such as silica creates a persistent oxidative stress-rich microenvironment in the lungs. The sustained oxidative stress promotes chronic lung inflammation and injury, AM death, and/or polarization toward pro-inflammatory (M1) or pro-fibrotic (M2) phenotypes, increased AT2 cell senescence and apoptosis, activation and expansion of lung fibroblasts, and ultimately PF^41,42,43^. Following phagocytosis by AMs, silica particles trigger the generation of ROS, including hydrogen peroxide, superoxide, singlet oxygen, and hydroxyl radicals. Recent studies provide further evidence that targeting silica-induced ROS in AMs could attenuate profibrotic signaling and fibrosis remodeling^42,43^. To this end, we first investigated the effects of CSP7 on silica-induced ROS generation, mice were exposed to varying concentrations of silica for different time. Later, lung tissues were analyzed for ROS generation. As shown in **Figure 3A**, *i.t.* administration of silica induced a concentration-and time-dependent increase in ROS generation. Conversely, treatment with CSP7 caused a significant reduction in ROS generation, as evidenced by marked reduction in ROS-associated fluorescence intensity. Further analysis of MMP revealed a concentration-dependent decline in MMP following silica exposure for 30 minutes to 4h (**Figure 3B)**, suggesting silica-induced mitochondrial dysfunction. Treatment with CSP7 markedly restored MMP, suggesting that its protective effects in resolving sPF may involve, at least in part, restoration of MMP. These findings were further corroborated by immunofluorescence (IF) analysis, which confirmed the restoration of MMP with CSP7 treatment. Furthermore, MitoSOX analysis indicated a concentration-dependent increase in following silica exposure in WT mice with sPF (**Figure 3C**). Notably, CSP7 treatment significantly reduced mitochondrial superoxide generation in the sPF+CSP7 group compared with untreated sPF group. Collectively, these findings further substantiate the multi-mechanistic therapeutics effects of CSP7 in mitigating silica-induced lung injury and fibrosis.

**Figure 3.**
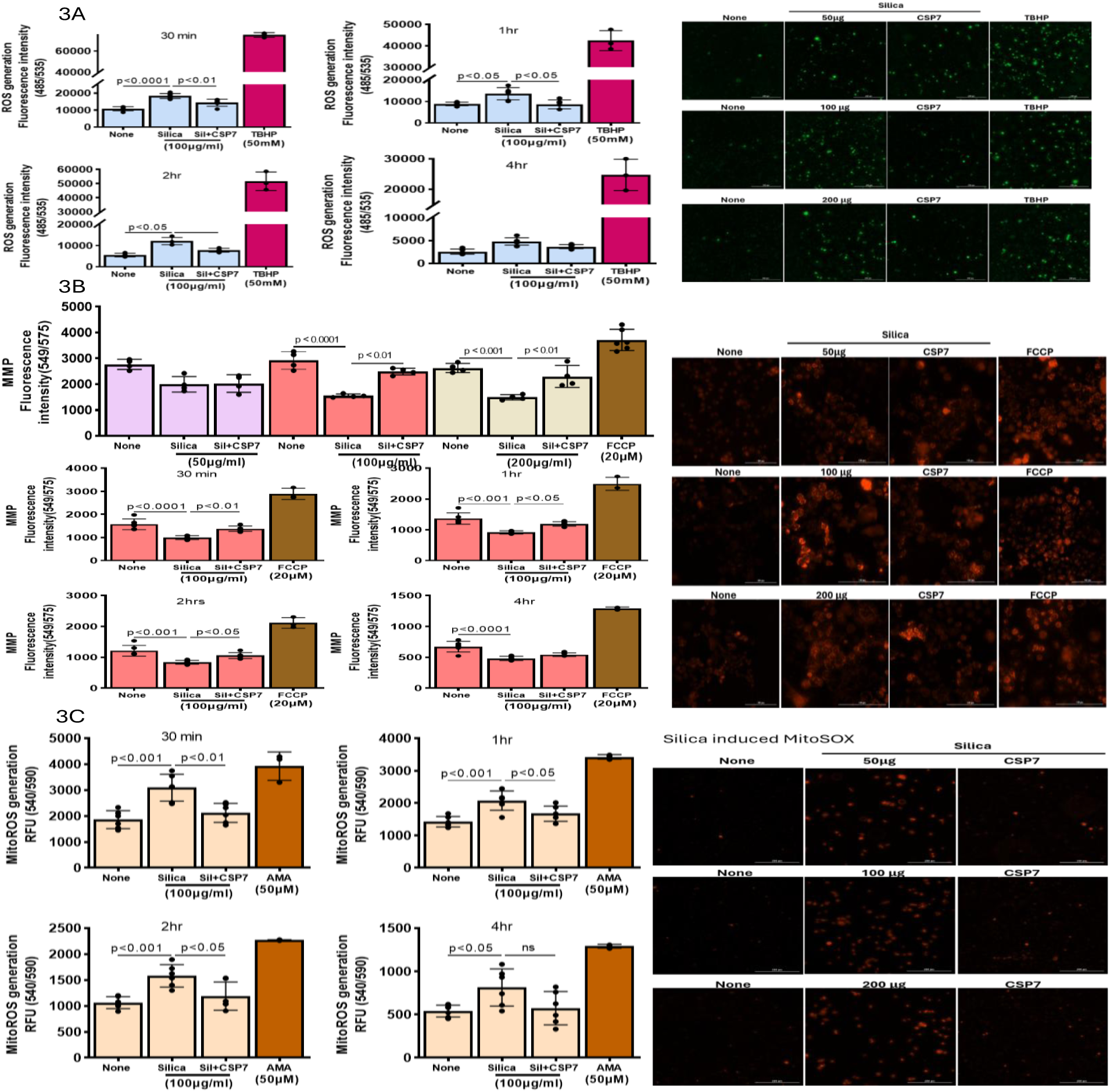
CSP7 inhibits ROS and MitoSOX production and restores MMP during silica induced lung injury. (**A**) CSP7 inhibits ROS production in WT mice with silica induced lung injury. The representative IF image of ROS induction in WT mice using different concentrations of silica (50, 100 and 200 ug/mL) for different time (0.5-4h) with or without CSP7. (**B**) **CSP7 restores MMP after silica induced injury.** The representative IF image of MMP induction in WT mice treated with silica and CSP7 (**C**) **CSP7 inhibits silica induced MitoSOX in WT mice.** The representative IF image of MitoSOX induction in WT mice exposed to silica with or without CSP7.

### CSP5 recapitulates the anti-apoptotic activities of CSP7 in human AT2 cells exposed to silica *in vitro* and mitigates sPF in WT mice

Our previous studies demonstrated that CSP and CSP7, both containing the CSP5 sequence, inhibit silica-induced apoptosis in AT2 cells. To determine whether CSP5 retains cytoprotective activity, human AT2 cells subjected to silica-induced injury were treated with CSP5 (20 µM) for 48h. Subsequently, AT2 cell lysates were analyzed for apoptosis markers (as indicated by the ratio of Cl. Cas-3/Cas-3) along with the expression of PAI-1 and SP-C by Western blotting technique. As shown in **Figure 4A**, exposure of AT2 cells to crystalline silica particles augmented activation of Cas-3 reflected by increased Cl. Cas and PAI-1, accompanied by reduced SP-C expression. These changes were reversed following treatment of AT2 cells with CSP5 initiated 24h after silica exposure, consistent with recovery of AT2 cell homeostasis.

**Figure 4.**
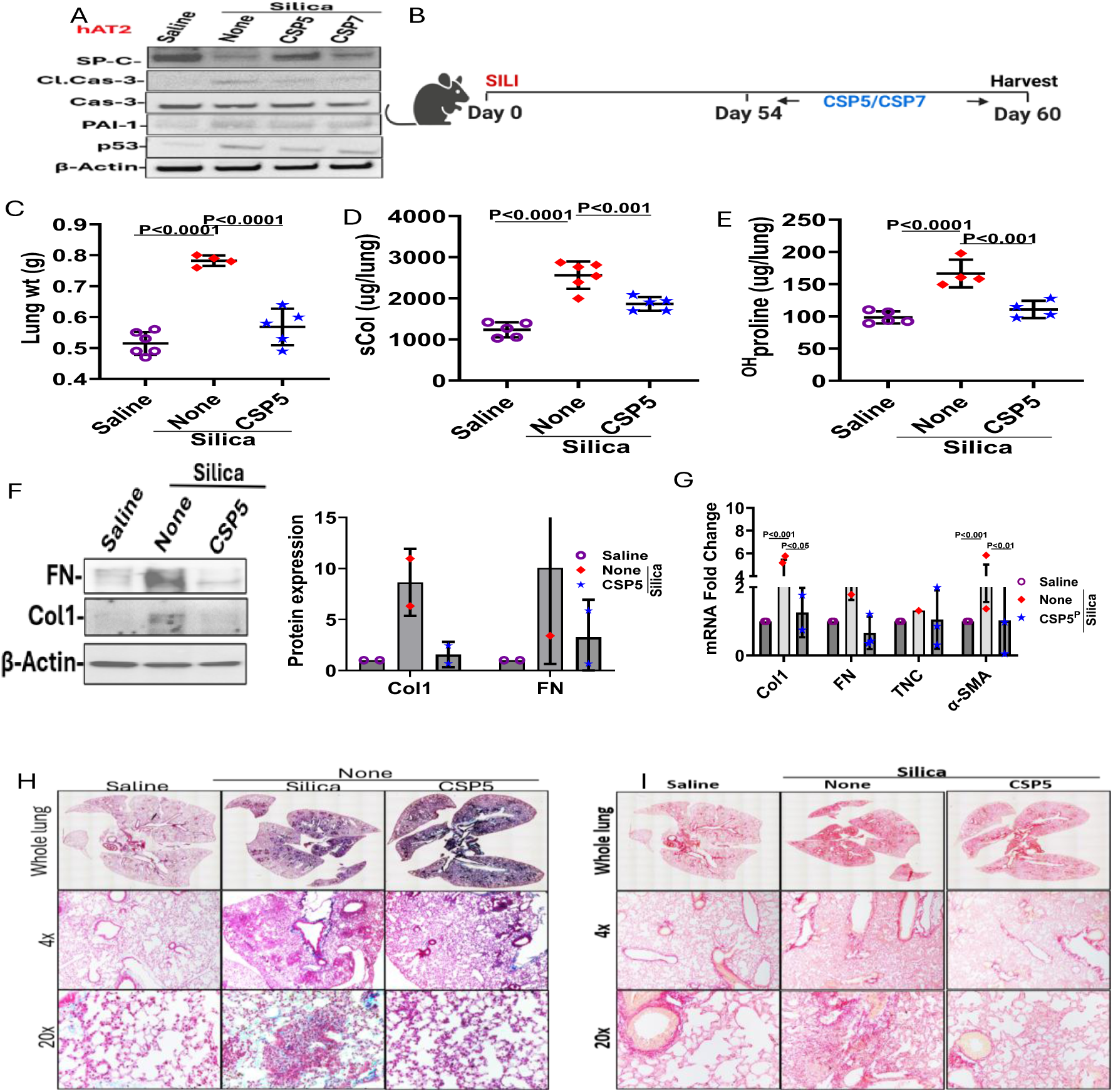
CSP5 mitigates silica induced injury in human AT2 cells *in vitro* and reverses PF in WT mice exposed to Silica. **(A**) Human AT2 cells were exposed to silica (100 mg/ml). After 24h, these cells were treated with 10 mM CSP5 or CSP7 for additional 48h. AT2 cell lysates were analyzed for SP-C, Cl. Cas-3, PAI-1 and b-actin by Western blotting. The bar graph represents densities of individual bands (n=3) after normalization with corresponding b-actin loading controls. (**B**) Schematic representation of experimental design. WT mice (n=5/gp) were exposed to silica for 60 days to induce Silicosis. After 54 days, mice exposed to silica were left untreated (None) or treated with CSP5 (1.5 mg/kg/day) by IP injection for 7 days. (**C**) Lung weight was measured 60 days after initiation of silica induced lung injury. Whole lung homogenates from naïve mice and mice exposed to silica and treated with or without CSP5 were analyzed for (**D**) sCol, (**E**) Total ^OH^Proline contents, (**F**) Profibrotic markers Col1 and FN proteins by Western blot. The densities of individual bands (n=3) were presented as bar graph. (**G**) Total lung RNA was subjected to qPCR for fibrotic markers mRNA. Lung sections were subjected to (**H**) Trichrome and (**I**) Picrosirius staining.

To further evaluate the therapeutic effects of CSP5 in mice with established sPF, WT mice received a single *i.t.* instillation of silica (1.5 mg/100 μl). After 54 days of post-exposure to silica, mice with established sPF were treated with CSP5 (1.5 mg/kg body weight) once daily by *IP* injection for seven consecutive days (**Figure 4B**). Fibrosis-associated endpoints were assessed on day-60 following silica exposure. As shown in **Figure 4C**, silica exposure significantly increased lung weight compared with saline-treated controls, consistent with fibrosis-associated tissue remodeling and ECM protein accumulation. Importantly, CSP5 treatment significantly reduced lung weight in mice with sPF compared with untreated sPF controls, suggesting attenuation of fibrotic remodeling. This reduction in lung fibrosis burden was further supported by biochemical assessments of sCol and ^OH^Proline contents in whole-lung homogenates. As shown in **Figure 4D**, treatment with CSP5 significantly reduced sCol levels in lung tissues of mice with established sPF. Similarly, the total lung ^OH^Proline levels were markedly reduced following CSP5 treatment **(Figure 4E)**, indicating attenuation of collagen accumulation and lung remodeling. Consistent with these findings, immunoblotting of whole-lung homogenates for ECM proteins such as Col1 and FN1 (**Figure 4F**), revealed significant reductions with CSP5 treatment. Furthermore, qPCR analysis demonstrated significant downregulation of profibrotic transcripts, including Col1, FN1, TN-C or a-SMA in CSP5-treated mice compared with untreated sPF controls (**Figure 4G**). Together, these findings demonstrate that CSP5 effectively attenuates established sPF by suppressing profibrotic gene expression, collagen accumulation, and ECM deposition. In addition, histopathological evaluation of Masson’s trichrome (**Figure 4H**) and picrosirius (**Figure 4I**) stained lung sections revealed marked reduction in interstitial deposition of Col and other ECM proteins following CSP5 treatment in mice with established sPF. Collectively, these findings demonstrate the therapeutic efficacy of CSP5 in attenuating silica-induced lung injury and resultant sPF and support its potential development as a treatment for silicosis, and other occupational particulate-induced progressive fibrotic lung diseases, including pneumoconiosis. Importantly, CSP5 treatment was initiated at an advanced stage of disease, after PF had already developed, demonstrating its ability to mitigate established fibrotic pathology rather than merely preventing the initial injury or onset of fibrosis.

### Role of autophagy, ER stress and AT2 cell specific MCP1IP1 expression in 1X-BLM-PF, sPF and TSLI

Earlier studies from our group demonstrate elevated expression of p53, endoplasmic reticulum (ER) stress-associated proteins, and autophagy-related proteins in AT2 cells isolated from the lungs of WT mice with BLM-PF, sPF or TSLI^4,36^. These alterations were attenuated by conditional deletion of p53 in AT2 cells or treatment with CSP7, suggesting that the antifibrotic effects of CSP7 may be mediated, at least in part, through suppression of p53 signaling, ER stress, and dysregulated autophagy. To test whether CSP5 affects selected autophagy-associated pathway proteins, we analyzed lung homogenates from mice with BLM-PF, sPF and TSLI for changes in autophagy proteins. Lung homogenates of mice with BLM-PF (**Figure 5A**), sPF (**Figure 5B**) and TSLI (**Figure 5C**) showed elevated expression of several autophagy-associated proteins, including p62, LC3B, Beclin-1, ARG1, and SOCS3, compared with the corresponding control groups of mice. Consistent with mitigation of BLM-PF, sPF or TSLI in mice, above mentioned autophagy related markers were remarkably down-regulated in the lung tissues of BLM-PF, sPF or TSLI mice treated with CSP5 or CSP7. Thus, indicating reversal of autophagy dysregulation may reverse lung injury and subsequent fibrotic remodeling. In addition, immunoblotting of lung homogenates of BLM-PF, sPF or TSLI mice revealed marked increase in monocyte chemotactic protein-induced protein 1 (MCPIP1 aka Regnase-1), which was reduced in mice exposed CSP5 or CSP7. Previous studies have implicated MCPIP1 in regulation of ER stress responses by influencing mRNA stability, stress granule dynamics, and autophagy, thereby affecting cellular stress responses and survival. Further MCPIP1 is involved in p53-associated autophagy signaling, suggesting a mechanistic link in p53-mediated induction of MCPIP1 with PF. Together, these findings suggest an interplay among inflammatory signaling, ER stress, p53 signaling, and dysregulated autophagy in AT2 cells, potentially contributing to alveolar epithelial injury and the subsequent development of PF.

**Figure 5.**
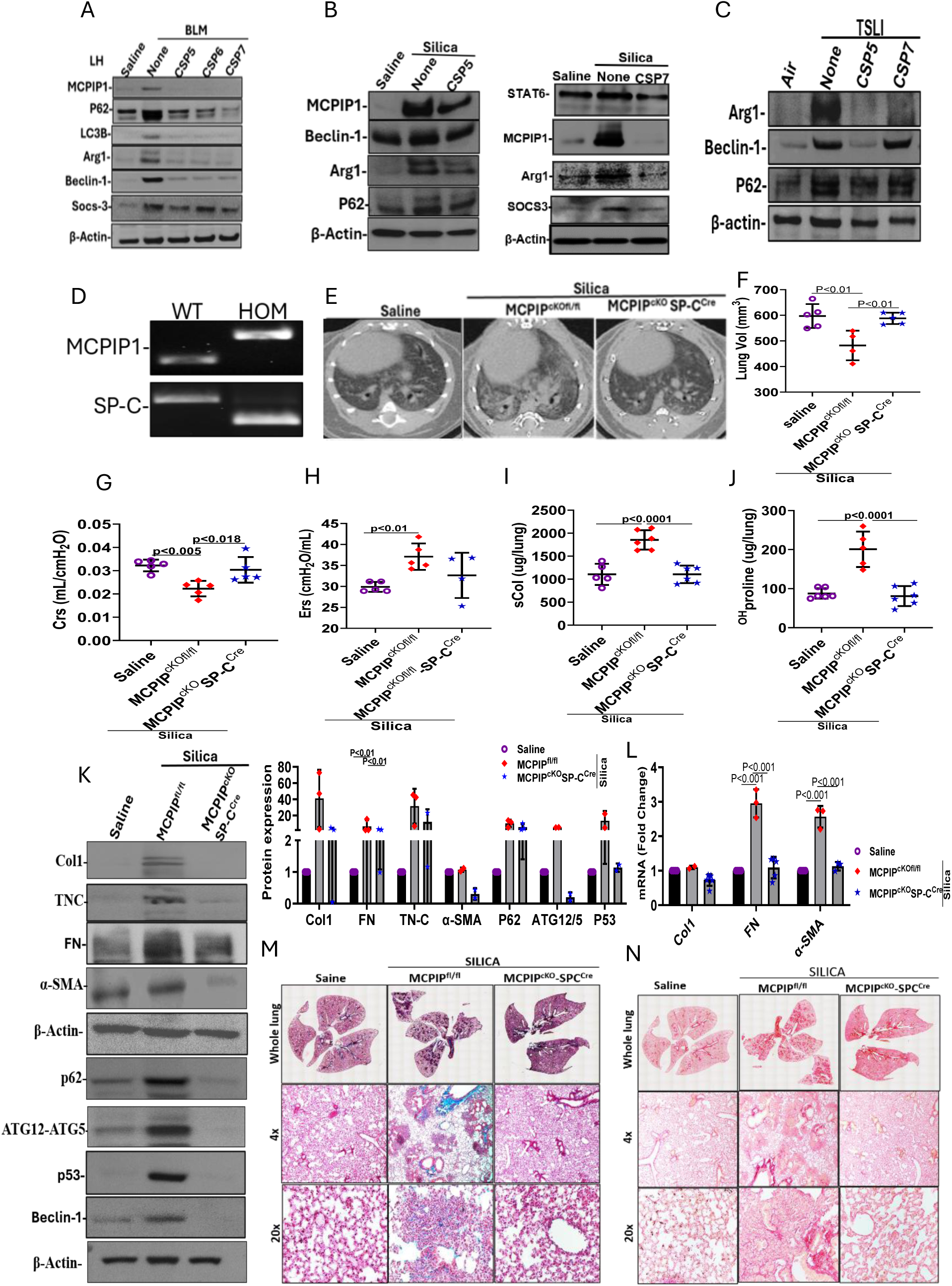
AT2 specific MCPIP1^cKO^ mice mitigate sLI and PF. Lung homogenates of WT mice with (**A**) 1X-BLM-PF, (**B**) sPF and (**C**) TSLI were treated with CSP5, CSP6 and CSP7 as described in Figures 1, 2 and 4 were analyzed for Autophagy markers and MCPIP1 using western blot. (**D**) Generation of AT2 cell specific MCPIP1^cKO^ mice by breeding MCPIP1^fl/fl^ and SP-C^Cre^ mice. The mice were IP injected with tamoxifen to induce Cre recombinase and exposed to saline or silica by *i.t.* instillation to induce sPF as in Figure 4B. Sixty days after saline and silica treated MCPIP1^fl/fl^ and MCPIP^cKO^ mice (n=5/gp) were subjected to (**E**) μCT imaging, and (**F**) Lung volume after μCT rendition, (**G**) Lung compliance Crs and (**H**) Lung elastance Ers were measured from these mice. Whole lung homogenates were analyzed for (**I**) sCol contents, (J) Total ^HO^Proline and (**K**) Profibrogenic markers and autophagy pathway proteins by Western blotting. The densities of individual bands (n=3) were presented as bar graph after normalization with corresponding b-actin loading controls. (**L**)Total lung RNA was analyzed for Col1, FN, TN-C and a-SMA mRNA by qPCR analyses. Lung sections from these mice were subjected to (**M**) Masson’s Trichrome and (**N**) Picrosirius staining.

To investigate the contribution of AT2 cell-specific MCPIP1 expression in the development of sPF and BLM-PF, we generated tamoxifen-inducible, AT2 cell-specific conditional knockout (MCPIP1^cKO^) mice (**Figure 5D**). As shown in **Figure 5E**, µCT scanning revealed markedly reduced fibrotic scarring in the lungs of MCPIP1^cKO^ mice compared with MCPIP1^fl/fl^ controls following silica exposure. Further, lung volume (**Figure 5F**), Crs (**Figure 5G**), and Ers (**Figure 5H**) were substantially improved in silica treated MCPIP1^cKO^ mice compared with MCPIP1^fl/fl^ mice. Further biochemical analyses of whole-lung homogenates revealed significantly lower sCol (**Figure 5I**) or total ^OH^Proline contents (**Figure 5J**) in silica-exposed MCPIP1^cKO^ mice. Consistent with these findings, Western blot analysis revealed remarkably reduced expression of the profibrotic proteins namely, Col, FN1, TN-C and a-SMA as along with autophagy-associated proteins p62 and the ATG12–ATG5 complex, in the silica-exposed MCPIP1^cKO^ mice relative to MCPIP1^fl/fl^ controls (**Figure 5K**). Complementing these findings, qPCR analysis of whole-lung homogenates demonstrated significant suppression of Col1, FN1, TN-C or a-SMA mRNA (**Figure 5L**) in MCPIP1^cKO^ mice exposed to silica compared with silica-exposed MCPIP1^fl/fl^ controls. Additionally, histopathological evaluation of Masson’s trichrome-(**Figure 5M**) and picrosirius red-stained (**Figure 5N**) lung sections confirmed markedly reduced interstitial collagen deposition and accumulation of other ECM proteins in AT2 cell-specific MCPIP1^cKO^ mice. Overall, these findings implicate AT2-cell specific MCPIP1 as a key contributor to silica-induced epithelial cell injury and subsequent sPF, potentially through modulation of ER stress and dysregulated autophagy-associated signaling pathways.

To further establish the contribution of AT2 cell-specific MCPIP1 expression to the development of PF, MCPIP1^cKO^ and MCPIP1^fl/fl^ mice were subjected to *i.t.* BLM (1U/Kg body weight). Evidently, tamoxifen-inducible, AT2 cell-specific MCPIP1^cKO^ mice exhibited marked protection to 1X-BLM-PF, reflected by improved overall survival (**Figure 6A**) and significantly attenuated body weight loss (**Figure 6B**) compared with BLM-exposed MCPIP^fl/fl^ controls, relative to saline-treated naïve mice. The resistance of MCPIP1^cKO^ mice to develop BLM-PF was further supported by a minimal increase in gross lung weight compared with marked increase observed in BLM-exposed MCPIP^fl/fl^ controls (**Figure 6C**). Additionally, biochemical analysis of whole-lung homogenates for total ^OH^Proline (**Figure 6D**) or sCol (**Figure 6E**) contents showed a significantly lower accumulation of ECM proteins in MCPIP1^cKO^ mice. The resistance of MCPIP1^cKO^ mice to develop BLM-PF was further confirmed by minimal increase in the expression of profibrogenic markers such as Col1, FN1, TN-C and a-SMA proteins by Western blot analysis (**Figure 6F**) and their corresponding transcripts by qPCR (**Figure 6G**) processed from whole-lung homogenates. Correspondingly, histopathological analysis of lung sections by Masson’s Trichrome (**Figure 6H**) and Picrosirius red staining (**Figure 6I**) revealed minimal deposition of Col and other ECM proteins, accompanied by preservation of alveolar architecture in 1X-BLM-exposed MCPIP1^cKO^ mice, in marked contrast to extensive fibrotic remodeling and architectural distortion observed in similarly treated MCPIP1^fl/fl^ controls. Taken together, these findings implicate elevated MCPIP1 expression in AT2 cells pivotal to BLM-induced AT2 cell damage and subsequent development of PF, processes that are attenuated by AT2 cell-specific MCPIP1 deficiency or treatment with CSP5 or CSP7, which share the CSP5 peptide sequence and suppress MCPIP1 expression.

**Figure 6.**
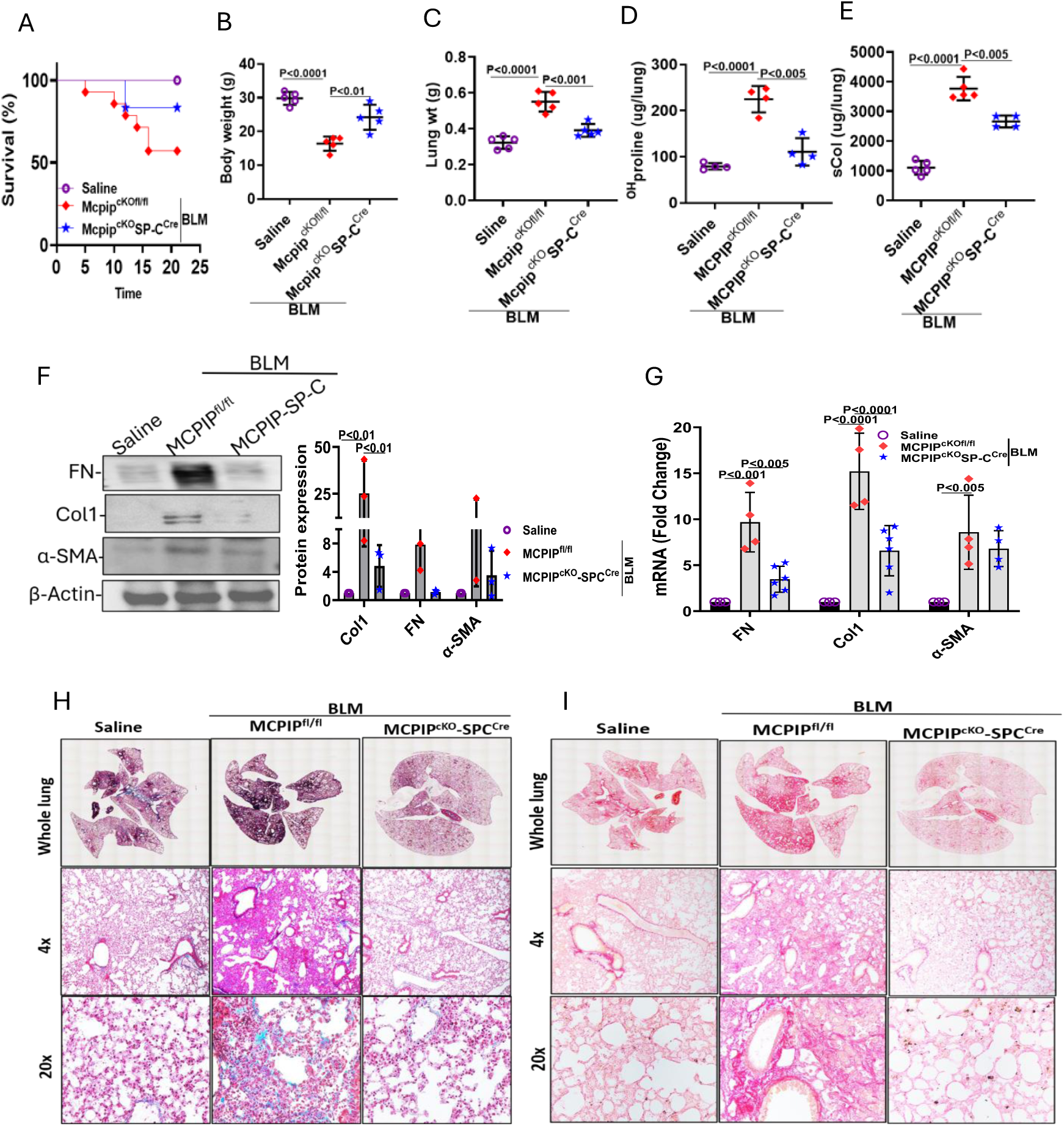
AT2-specific MCPIP1 deficiency protects mice from developing BLM-PF. MCPIP1^fl/fl^ and MCPIP1^cKO^ mice (n=5/gp) were exposed to 1X-BLM days to induce PF. Mice were evaluated for post-BLM (**A**) Survival during 21 days. The changes in (**B**) Body weights and (**C**) Lung weights were noted twenty-one days after initiation of BLM injury. Whole lung homogenates were analyzed for (**D**) Total ^OH^Proline and (**E**) sCol contents, and profibrogenic (**F**) Western blotting for pro-fibrogenic marker proteins. The densities of individual bands (n=3) are presented as bar graph after normalization with corresponding b-actin loading controls. Whole lung homogenates were analyzed for profibrogenic markers mRNA by qPCR (**G**). Lung sections were subjected to (**H**) Masson’s Trichrome and (**I**) Picrosirius red staining for Col and other ECM protein deposits *in situ*.

MCPIP1 has been shown to regulate autophagy and promote alternative polarization of Mo-AMs to pro-fibrotic M2 phenotypes, potentially contributing to fibrotic remodeling^44^. In fibrotic lungs, including mouse PF lungs and human IPF lungs, alternatively programmed (M2) Mo-AMs are elevated, and they induce fLf activation and impair alveolar epithelial regeneration through elaboration of TGF-b and other profibrotic cytokines^45^. Therefore, we sought to determine whether antifibrotic activity of CSP5 or CSP7 encompassing the CSP5 sequence, involves suppression or reversal of profibrotic M2-like polarization of Mo-AMs. As shown in **Figure 7A** and **7B**, flow cytometry analysis of AMs isolated from BLM-PF mice demonstrated a marked increase in the proportion of CD206 and CD71 positive cells, with minimal changes in CD80 positive population, compared with AMs isolated from saline-treated naïve control mice, suggesting increased accumulation M2-like macrophages in fibrotic lungs. This was significantly reduced following treatment of BLM-PF mice with CSP7. Interestingly, CSP7 did not increase CD80 population, indicating CSP7 treatment inhibits M2 polarized AMs without increasing pro-inflammatory M1 phenotype. We next analyzed AM lysates for expression for M2 markers (MCPIP1, KLF4, Stat6, SOCS3, Arg1 and CD206), autophagy markers (LC3BI, LC3BII, p62 and ATG12-ATG5), Cav1, profibrotic cytokines such as TGF-b1 and CTGF (**Figure 7C**) and ER stress markers (BiP, GRp78 and CHOP) (**Figure 7D**) by Western blotting, qPCR for MCPIP1, SOCS3, Arg1, MIP1a, CD206, TGF-b1 and CTGF mRNA (**Figure 7E**) and IHC and IF imaging of MCPIP1, LCB and Arg1 (**Figure 7F**). Consistent with flow cytometry findings, we found marked decrease in markers of M2 phenotypes and autophagy along with inhibition of MCPIP1 without significantly affecting MIP1a expression in Mo-AMs isolated from BLM-PF mice treated with CSP7 compared to their corresponding levels in BLM-PF mice left untreated. Thus, collectively confirming that CSP7 inhibits profibrotic polarization of Mo-AM by affecting autophagy and ER stress pathways.

**Figure 7.**
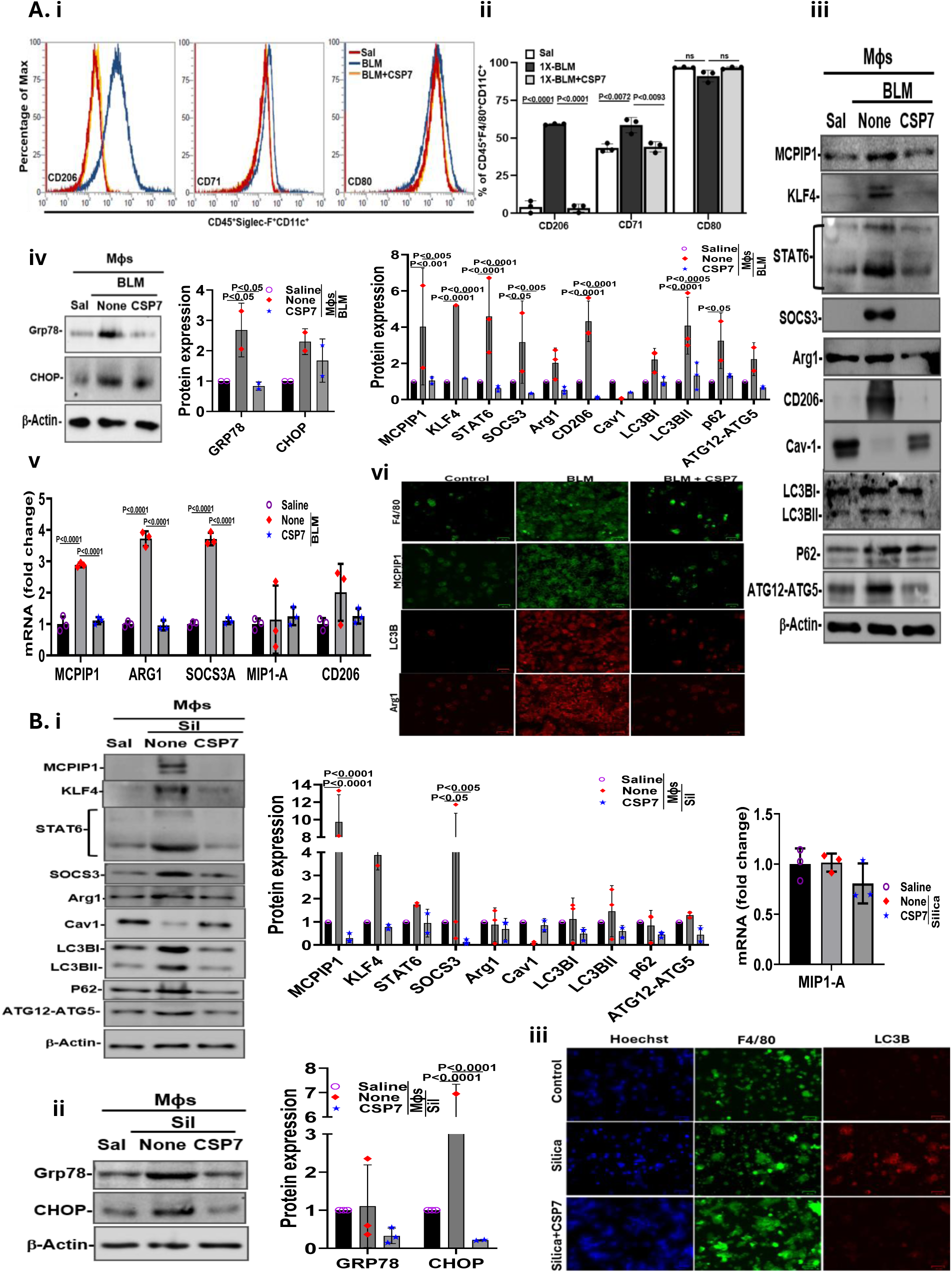
**CSP7 Mitigates the BLM and silica induced autophagy dysregulation and ER stress in AMs.**(**A**) WT mice (n=5/gp) were exposed to 1X-BLM for 21 days to induce PF. After 14 days, mice exposed to BLM were left untreated (None) or treated with CSP7 (1.5 mg/kg/day) by IP injection for 7 days. (**i**) Flow cytometry analysis and (**ii)** Quantification of macrophage polarization. (**iii**) Lysates of AMs were immunobloted for MCPIP1, Cav1 and autophagy and (**iv**) ER stress markers. (**v**) Total RNA isolated from AMs were analyzed for MCPIP1, SOCS3 and Argq mRNA by qPCR. (**vi**) IF images of autophagy markers in isolated AMs. (**B**) WT mice (n=5/gp) were exposed to Silica to induce sPF and 54 days after i.t. instillation of silica, mice with sPF were left untreated (None) or treated with CSP7 (1.5 mg/kg/day) by IP injection for last 7 days from day 54 to 60. Lysates of AMs were (**i**) Immunobloted for MCPIP1, Cav1 and autophagy markers and (**ii**) ER stress markers. (**iii**) AMs isolated sPF mice were subjected to IF staining for autophagy and macrophage markers.

Given the well-established role of AMs as the first line of defense against inhaled particulates and their critical role in the pathogenesis of sPF^35^. We further investigated the effect of CSP7 on AM polarization, autophagy and ER stress. To this end, we isolated AMs from naïve mice and sPF mice treated with or without CSP7. Cell lysates from AM were analyzed for M2 markers (MCPIP1, KLF4, Stat6, SOCS3 and Arg1) autophagy markers (LC3BI, LC3BII, p62 and ATG12-ATG5) and Cav1 (**Figure 7G**) as well as for ER stress markers (GRp78 and CHOP) (**Figure 7H**) by Western blotting. Additionally, IHC and IF analysis were performed to evaluate the expression of LC3B and F4/80 (**Figure 7I**). Immunoblot and qPCR findings showed that CSP7 significantly downregulated the silica-induced aberration of autophagy-associated markers in Mo-AMs. IF Images also confirmed a notable decrease in the fluorescence intensity of LC3B in AMs after CSP7 treatment when compared to their fluorescence intensity in AMs isolated from sPF mice, corroborating that CSP7 can mitigate silica-induced changes in autophagy-associated proteins. We next explored the role of CSP7 in the regulation of ER stress in AMs of sPF mice. Our results demonstrated increased the expression of the ER stress-associated proteins GRP78 and CHOP in Mo-AMs of sPF mice, which is reduced after CSP7 treatment. These findings collectively demonstrate that CSP7 mitigates silica-induced autophagy pathway dysregulation and ER stress in AMs.

## Discussion

CLDs, including IPF, silicosis, COPD, and several other lung disorders are debilitating, and progressive conditions lacking curative therapies. Current treatment strategies vary by disease type but primarily focus on providing symptomatic relief, slow down progression, reduce inflammation, improve airflow, and prevent exacerbations, rather than reverse disease pathology and restore lung architecture. While current therapies, including bronchodilators, inhaled corticosteroids, selective phosphodiesterase inhibitors, and biologics can reduce exacerbations and improve lung function and quality of life, their ability to substantially modify the underlying pathophysiology of the disease remains limited. Smoking cessation, long-term oxygen therapy in selective patients, lung-volume reduction procedures, and lung transplantation have demonstrated to prolong life in specific patient populations. Nevertheless, pharmacological research in CLD has largely focused on targeting individual molecular mediators, signaling pathways, or a specific cell population despite growing evidence clearly indicates that the underlying pathobiology involves complex interaction of multiple cell types and interconnected molecular pathways. The continued absence of curative therapies for CLDs underscores the critical need for identifying pharmacological targets that concurrently modulate multiple cellular processes and interconnected disease-driving pathways, thereby enabling resolution of lung pathology and restoration of pulmonary architecture and function.

Caveolins are structural proteins of plasma membrane vesicular invaginations referred to as caveolae. Cav1 dynamically traffics between plasma membrane, intracellular organelles, and the cytoplasm^14–18^, where it functions as a scaffolding protein organizing signaling complexes. We and others have demonstrated elevated expression of Cav1 in AT2 cells and AE cells from patients with COPD and mice subjected to TSLI, as well as AT2 cells from patients with IPF and mice with BLM-PF and sPF. The process is intricately linked to increased expression of p53 and PAI-1 in these cells^46–48^. In addition, overexpression of Cav1 promotes induction of p53 and p53-mediated downstream induction of PAI-1 and suppression of uPA and uPAR expression. This in turn induces senescence and apoptosis resulting in reduced AT2 cell viability, and MCM and MH by AE cells. The coordinated changes of decreased uPA and uPAR with increased PAI-1 are consequent to induction of p53 that occurs in injured AT2 and AE cells *via* Cav1-mediated signaling^6,48–53^. These findings highlight the regulatory crosstalk between p53 and uPA system, whereby p53 binds to component mRNA 3′ UTR sequences^48–53^. Together, these studies implicate elevated Cav1 expression in AT2 cell senescence, apoptosis, and loss of alveolar structural integrity, and activation and expansion of fLfs, providing a compelling predicate for targeting Cav1-mediated signaling pathways as a therapeutic strategy for both obstructive and restrictive lung diseases. In addition, we and others have shown that Cav1 scaffolding domain 20-mer peptide CSP inhibited p53 in AT2 cells, which increased AT2 viability and attenuated PF. Furthermore, we demonstrated that CSP7, a 7-mer deletion fragment of CSP, modulates multiple signaling pathways governing cellular viability in AT2 cells, activation and differentiation of fibroblasts, and AM fate, polarization, and function, thereby mitigating lung injury and PF at various stages of disease progression^4,6,22,48,49^. In addition, Cav1 interacts with both autophagy (LC3 and ATG12-ATG5)^36,54,55^ and apoptotic proteins (Fas and BID), thus linking extrinsic and intrinsic apoptotic pathways^56^. Growing evidence from patients with IPF and experimental models of PF suggest that dysregulated autophagy promotes AT2 cell injury and apoptosis while facilitating fLf activation and differentiation, ultimately leading to excessive deposition of ECM. We further demonstrated that CSP or CSP7 delivered by *IP* injection after solubilization in buffer containing DMSO, or micronized CSP7 without excipients resolves established PF in multiple preclinical models and mitigates TSLI^4,6,21,36^. In contrast, CSP and its 7-amino acid derivative, CSP7 without penetrating sequences, were developed in our laboratory to target Cav1-dependent cellular alterations, preserve AT2 cell homeostasis, and attenuate lung injury and the progression of PF. CSP7 peptide, administered *via Neb*, *DPI*, or *IP* injection, attenuates lung injury and PF while improving lung function across multiple preclinical mouse models^4,6,21,27,36^. Toxicity studies of micronized CSP7 (aka LTI-03) in mice, rats, dogs, along with Phase 1a safety and pharmacokinetic (PK) studies in healthy human subjects (NCT04233814)^28^ and Phase 1b safety studies in patients with newly diagnosed IPF (NCT05954988)^29,30^, have been completed. Currently, CSP7 has advanced to a multicenter Phase 2 (NCT06968845)^31^ clinical trial evaluating its therapeutic efficacy in patients with IPF across clinical sites in the US and Europe.

However, administration of the poorly water-soluble CSP7 necessitates micronization^21^ or dry powder formulation by thin film freezing^27^ for inhalation, and requiring advanced procedures and specialized equipment’s to deliver the medication. Furthermore, regardless of the *DPI* formulation, the deposited peptide particles must dissolve in lung lining fluid to facilitate drug availability and subsequent pharmacological activity. The limited aqueous solubility of micronized CSP7 (LTI-03) powder may therefore present challenges for efficient drug delivery and long-term tolerability. Notably, minimal bronchiolar MC hyperplasia was reported in the 26-week rat inhalation study and specifically highlighted by FDA^34,35^. This finding raises particular concern regarding the development of CSP7 for COPD, a disease characterized by goblet cell metaplasia/ hyperplasia, MH, impaired MCC and small airway obstruction. To address these limitations, we investigated whether truncated derivatives of CSP7 could retain or enhance the biological activity of the parent peptide while exhibiting improved physicochemical properties, particularly greater aqueous solubility, with a potential to minimize local epithelial cell irritation and overcome formulation-and dissolution-related challenges associated with CSP7. In the present study, we evaluated the relative therapeutic potency of CSP5 and CSP6, the two truncated derivates of CSP7, against BLM-PF, sPF and TSLI in established *in vitro* and *in vivo* models. Notably, among the three peptides evaluated, CSP5 exhibited superior physicochemical properties, especially greater aqueous solubility, along with enhanced AT2 cell viability. Compared with its parent peptide, CSP7, CSP5 exhibited remarkable efficacy in attenuating the expression of apoptotic markers, including p53, PAI-1 and Cl. Cas-3, while promoting the restoration of SP-C expression by AT2 cells. Interestingly, despite the differences in physicochemical properties and *in vitro* biological activity, CSP5, CSP6, and CSP7 exhibited antifibrotic activity across *in vivo* experimental models. All three peptides demonstrated substantial improvements in pulmonary function and lung architecture, as evidenced by μCT imaging, lung function assessments, histopathological examination, and morphometry analyses. These findings were further corroborated by reductions in ^OH^Proline and sCol content, along with molecular changes revealed by western blotting and qPCR. Together, these findings demonstrate that the truncated derivative of CSP7, CSP5 retain antifibrotic efficacy comparable to the parent peptide. Although all three peptides demonstrated beneficial therapeutic activity across the preclinical models, CSP5 emerged as a particularly promising candidate for further development owing to its greater aqueous solubility, enhanced potency to preserve epithelial cell viability and regeneration capacity, and robust antifibrotic activities, consistent with previously reported efficacy of CSP7 towards the protection of AT2 cells through suppression of p53 and PAI-1 pathway^6^. These distinguishing features of CSP5 support its continued development as a potential therapeutic for CLDs.

Autophagy is an important mechanism that maintains cellular homeostasis by regulating the recovery of damaged cellular components and organelles. However, repeated lung damage can dramatically alter the autophagy pathway, resulting in epithelial dysfunction, oxidative stress, inflammation, and fibrotic remodeling^54^. In the present study, BLM-PF, sPF and TSLI were accompanied by notable alterations in autophagy-associated markers, including p62, LC3B, and Beclin-1^36^. Interestingly, treatment with CSP5 and CSP7, which encompasses the CSP5 sequence, significantly reduced the expression of these proteins. These findings are consistent with our previous reports linking the attenuation of PF and TSLI to modulation of the autophagy-related pathways. Dysregulated autophagy impairs the clearance of damaged cellular components, exacerbate cellular stress, and inflammatory signaling, consequently favoring epithelial cell damage, fibroblast activation, and excessive ECM deposition. Together, these findings suggest that modulation of autophagy-related pathways may contribute to the protective effects of CSP5 and CSP7. Furthermore, treatment with CSP5 and CSP7 peptides significantly reduced the expression of ARG1 and SOCS3 in mice with BLM-PF and sPF. ARG1 is associated with AM activation and profibrotic responses, whereas SOCS3 is a key regulator of cytokine signaling that modulates inflammatory responses through JAK/STAT signaling pathway^36,57^. These findings suggest that CSP5 and CSP7 may attenuate AM-associated inflammatory signaling by modulating the ARG1- and SOCS3-associated pathways. Together, the present study demonstrates that CSP5, a truncated derivative of CSP7, effectively reduces the accumulation of ECM proteins and protects AT2 cells against apoptosis across BLM-PF, sPF, and TSLI models.

TS exposure is a well-established risk factor for CLDs, and increasingly recognized as a significant contributor to the pathogenesis of IPF, and may exacerbate sPF. Especially, an imbalance in oxidant-antioxidant ratio, excess mucus secretion, telomere dysfunction (TD) along with AT2 cell damage promoting senescence, apoptosis, necrosis, accompanied by accumulation of oxidative DNA damage and lipid peroxidation intermediates, are characteristics of TS-induced lung inflammation and injury^58,59^. Furthermore, TSLI destroys the lung tissues, contributing to parenchymal scarring. In addition to contribution to the progression of PF, TSLI accelerates the development of emphysema in individuals with preexisting emphysema. Therefore, patients with CPFE often exhibit relatively preserved lung volumes and FVC, despite progressive lung damage^60^. The impairment in epithelial regeneration contributing to progressive alveolar wall destruction and emphysema in TSLI are intricately linked to induction of p53, PAI-1, TD, senescence/apoptosis in AT2 cells^4,6^. In addition to promoting telomere shortening, TD, and reduced viability of AT2 cells, TSLI induces MCM and M5Ac in AE cells, partly through epigenetic suppression of forkhead box A2, causing impaired MCC, and airway remodeling along with mucus plugging^4,61–63^. We further demonstrated that p53-induced PAI-1 expression promotes MCM, and MH in AE cells, thereby contributing to the pathogenesis of TSLI. These pathological alterations were reversed by CSP7 encompassing the CSP5 sequence^4^, supporting its therapeutic potential for both chronic bronchitis and emphysema associated with COPD.

Consistent with these histologic findings, molecular analyses demonstrated reduced airway mucus markers following CSP5 treatment while restoring expression of SP-C. Corresponding reductions in M5Ac and restoration of SP-C transcripts relative to untreated TSLI mice. These findings indicate preservation of alveolar epithelial integrity and support a protective effect of CSP5 on alveolar repair mechanisms. These findings demonstrate that CSP5 improves airway and alveolar pathology and reduces TSLI-associated lung enlargement, improved lung function, reduced goblet-cell metaplasia and MH, decreased M5Ac, and restored SP-C expression in lung tissue and isolated AT2 cells. Surprisingly, CSP5 produced therapeutic effects comparable or better than those observed with CSP7 despite comprising a shorter peptide sequence. CSP5 additionally exhibits substantially improved aqueous solubility relative to CSP7. The combination of comparable or better biological efficacy and improved physicochemical properties supports the use of CSP5 for chronic lung administration, including administration by *DPI*, *Neb*, or other inhaled or systemic delivery modalities. These findings provide a strong rationale for advancement of CSP5 as a therapeutic candidate for treatment of COPD, chronic bronchitis, emphysema, TSLI, CPFE and related lung diseases.

Notably, the present study demonstrates that CSP5 restores AT2 cell viability and SP-C expression to levels observed in uninjured control AT2s cells, attenuates alveolar structural destruction, and suppresses MCM through downregulation of M5Ac expression in *in vitro* and *in vivo* models of TSLI. The distinct phenotype of CPFE is the outcome of chronic lung injury and frequently observed in smokers^39^. In this context, CSP5 treatment significantly reduced the expression of fibrotic markers and attenuated the ECM deposition surrounding the major airways in TSLI mice. Collectively, these findings support the therapeutic potential of CSP5 in attenuating alveolar epithelial injury, mucus-associated airway pathology, and peribronchial fibrosis, thereby targeting multiple pathological features of COPD, including those associated with chronic bronchitis and emphysema.

Beyond its protective effects against TSLI, CSP5 also demonstrated therapeutic activity against sPF, characterized by persistent inflammation, alveolar epithelial injury, and progressive fibrotic remodeling. Notably, expression of the pro-inflammatory cytokine IL-17A, is significantly elevated in sPF lungs^38^. IL-17A induces MCPIP1, tumor suppressor protein p53, and PAI-1 in AT2 cells^64^. In the present study, we further demonstrated that CSP5 blocked apoptosis in AT2 cells exposed to silica *in vitro*, and in isolated AT2 cells from sPF mice. Dysregulated autophagy and associated apoptotic responses are apparent in AT2 cells, contributing to the pathogenesis of sPF^36,65,66^. We observed significant increases in sCol and ^OH^Proline content in lung homogenates from silica-exposed WT mice compared with naive control mice. Additionally, the expression of pro-fibrogenic markers Col1, αSMA, FN, and TN-C protein, and their corresponding mRNAs were markedly elevated in mice with sPF. Lung sections stained with trichrome and picrosirius red further revealed increased Col and other ECM protein deposition in lung tissue sections, confirming the development of sPF. Notably, systemic administration of CSP5 or CSP7, which encompasses CSP5 sequence, via once-daily *IP* injection for seven consecutive days (day 54 through 60 post-silica exposure), significantly reduced sCol and total ^OH^Proline content in whole lung homogenates, demonstrating antifibrotic activity against established sPF. CSP5 treatment also markedly reduced the proteins and mRNAs of pro-fibrogenic markers, indicating mitigation of sPF, further corroborated by Trichrome and picrosirius red staining of the lung sections, which revealed diminished collagen deposition and fibrotic remodeling. We observed significant increase in lipidation of LCBII, along with elevated expression of autophagy substrate and cargo receptor protein, p62/SQSTM1, ATG5 at both protein and mRNA levels in the lungs of mice with sPF. Importantly, administration of CSP or CSP7 reversed these changes. Similarly, AT2 cells isolated from sPF mice showed increased expression of LC3BII, p62 and ATG5, p53, PAI-1, accompanied by enhanced apoptotic. These features were reversed following treatment with CSP5 or CSP7 starting day 54 through 60 following post-silica exposure, suggesting that both peptides protect AT2 cells against apoptosis and resolve sPF, potentially by regulating autophagy. Consistent with these findings, our previous studies demonstrated that exposure to silica failed to induce LC3BII, p62 and ATG5, or apoptosis in AT2 cells from p53^cKO^ mice, in which CSP7, containing the TFTVT sequence, had minimal additional effects^36^. These findings clearly suggest a central role for p53 in silica-induced dysregulation of autophagy, potentially mediated through MCPIP1 induction in injured AT2 cells, and demonstrate that CSP7 or CSP5 restore AT2 cell homeostasis by supressing p53, MCPIP1, and PAI-1.

We next analyzed ROS levels in AT2 cells using cell-permeant reagent 2’,7’-dichlorofluorescin diacetate (DCFDA) and mitochondrial (mt.) superoxide indicator, mitoSOX. We also analyzed MMP (ΔΨm) using the cell-permeant fluorescent dye, TMRE. We found that silica caused a dose-dependent increase in ROS production in human AT2 cells treated *in vitro*. This was significantly reduced following treatment of these cells with CSP7. Similarly, AT2 cells isolated from WT mice with sPF showed increased ROS production *vs* naïve control mice. This was significantly reduced after treatment of WT mice having sPF with CSP7 for 7 days from day 54 to 60. AT2 cells isolated from WT mice with sLI also exhibited increased MitoSOX, which was significantly reduced following treatment of mice with CSP7. AT2 cells isolated from IL-17A-or PAI-1-deficient mice exposed to silica did not show a significant increase in ROS, and CSP7 treatment had minimal effect on these mice. We also found that silica exposure caused a significantly decreased MMP in AT2 cells treated *in vitro*, which was reversed following treatment of AT2 cells with CSP7. Similarly, MMP was significantly reduced in AT2 cells isolated from WT mice with sPF, which increased in those mice treated with CSP7. Silica failed to reduce MMP in p53- and PAI-1-deficient AT2 cells and CSP7 had no significant effect in these cells.

In accordance with previous reports implicating dysregulated autophagy in AT2 cells promoting apoptosis, and subsequently to the development of PF, we observed elevated MCPIP1 expression in mice with BLM-PF and Spf^68^, which was attenuated by treatment with CSP5 or CSP7. Previous studies have also reported induction of MCPIP1 and PAI-1 by pro-inflammatory cytokine, IL-17A^67^. These findings suggest an interplay between IL-17A, MCPIP1, p53, and PAI-1 signaling in autophagic dysregulation and pathogenesis of CLDs. In this context, we developed MCPIP1^fl/fl^ mice and tamoxifen inducible AT2 cell-specific MCPIP1^cKO^ mice, and observed that MCPIP1^cKO^ mice resisted BLM-PF and sPF unlike similarly treated WT or MCPIP1^fl/fl^ mice. Further analysis of lung homogenates revealed minimal changes in sCol, ^OH^Proline, and pro-fibrogenic marker proteins and their corresponding mRNAs as well as MitoSOX, and ROS levels in MCPIP1^cKO^ mice compared with marked alterations observed in WT or MCPIP1^fl/fl^ mice exposed to silica. In addition, AMs isolated from mice with sPF exhibited increased MCPIP1 protein and mRNA expression, as along with elevated autophagy-associated markers. These alterations were significantly attenuated following CSP7 treatment from days 54 to 60 in silica-exposed WT mice with BLM-PF or sPF. These findings were further supported by IHC analysis of isolated AMs. Additionally, consistent with IHC findings, immunofluorescence (IF) imaging of AMs isolated from lungs of BLM-PF or sPF mice, revealed increased expression of autophagy-associated markers, ER stress markers, and macrophage polarization markers. Western blotting and mRNA analyses revealed that CSP7 treatment markedly reduced the BLM-or silica-induced dysregulation of autophagy-associated markers in AMs. IF images also confirmed a significant downregulation in the fluorescence intensities of MCPIP1, LC3B, and ARG1 in AMs following CSP7 treatment compared with untreated BLM-PF or sPF mice, further supporting the ability of CSP7, which includes the CSP5 sequence, to attenuate aberrant alterations in autophagy-associated proteins induced by BLM. We next determined whether CSP7 also regulates ER stress in AMs isolated from BLM-PF and sPF lungs. Our findings demonstrated that the BLM-PF or sPF mice increased the expression of the ER stress-associated proteins, particularly GRP78 and CHOP, which were markedly reduced following CSP7 treatment. Given that AM polarization is a marked feature in fibrotic lungs, including human IPF lungs, we characterized AM phenotypic alterations following CSP7 treatment. Flow cytometric analysis of AMs for the expression of CD206, CD71, and CD80, revealed an increased expression of CD206 and CD71 in AMs isolated from mice with BLM-PF. Contrastingly, treatment with CSP7 significantly reduced the CD206 and CD71 expressions without affecting CD80, suggesting a selective attenuation of profibrotic polarization of AMs, consistent with the observed suppression of TGF-b and CTGF expression. Overall, these findings demonstrate that CSP7 modulates AM phenotype and function by affecting dysregulated autophagy pathway, ER stress, and associated profibrotic signaling, thereby contributing to its antifibrotic activity.

In summary, CSP5, a 5-mer (TFTVT) truncated derivative of CSP7, retains comparable or superior biological activity to its parent peptide across multiple established preclinical models of airway injury, alveolar epithelial injury, and PF, despite its substantially reduced peptide length. Notably, CSP5 exhibited in greater aqueous solubility while preserving potent cytoprotective and anti-fibrotic effects. The improved solubility of CSP5 may reduce peptide self-association and aggregation, simplify formulation development, facilitate aerosolization and dissolution following pulmonary deposition, and potentially improve airway tolerability, enabling its potential development for inhaled therapies delivered via *DPI* or *Neb* as well as systemic administration through *IV* or subcutaneous routes. In addition, the dominant features of CSP5, establish it as a formidable candidate for chronic maintenance therapy, particularly for COPD, where long-term treatment requires favorable physicochemical characteristics, sustained therapeutic efficacy, and pulmonary tolerability. Building on these promising findings, our future studies will focus on characterizing the PK profile, long-term safety, and formulation properties of CSP5 to advance its development as a next-generation therapeutic for chronic obstructive and fibrotic lung diseases.

